# Human Anti-Glycan Reactivity is Driven by the Selection of B cells Utilizing Private Antibody Gene Rearrangements that are Affinity Maturated in Germinal Centers

**DOI:** 10.1101/2024.10.15.618486

**Authors:** J. Stewart New, Christopher F. Fucile, Amanda R. Callahan, Julia N. Burke, Randall S. Davis, Wayne L. Duck, Alexander F. Rosenberg, John F. Kearney, R. Glenn King

## Abstract

The human antibody repertoire is broadly reactive with carbohydrate antigens represented in the universe of all living things, including both the host/self-as well as the commensal microflora-derived glycomes. Here we have used BCR receptor cloning and expression together with single-cell transcriptomics to analyze the B cell repertoire to the ubiquitous N-acetyl-D-glucosamine (GlcNAc) epitope in human cohorts and dissect the immune phylogeny of this predominant class of antibodies. We find that circulating anti-GlcNAc B cells exhibiting canonical BMem phenotypes emerge rapidly after birth and couple this observation with evidence for germinal center-dependent affinity maturation of carbohydrate-specific B cell receptors *in situ* during early childhood. Direct analysis of individual B cell clonotypes reveals they exhibit strikingly distinct fine-specificity profiles for palettes of GlcNAc containing moieties. These results suggest that a generalized exposure to complex environmental glycans drives the steady state anti-glycan repertoire.

## Introduction

Anti-carbohydrate antibodies (ACA) are present in all human sera and are considered part of the natural antibody (nAb) repertoire. Though predominantly of IgM and IgA isotypes, ACAs also include IgG subclasses.^1^ The emergence of these antibodies appears to result from environmental exposures, as IgM glycan-reactive antibodies are nearly absent in cord blood, but emerge within the first few years of life and further accumulate with age.^2–4^ Some carbohydrates are variably targeted by ACAs of different individuals; for example, whether a person has ACAs reactive with Blood Group antigens is determined by the individual’s blood type. However, other ACAs are invariably observed in all individuals, comprising what has been described as a universal human ACA architecture.^3,5,6^ These conserved ACA targets include monosaccharides such as N-acetyl-Glucosamine (GlcNAc) and N-acetyl-Galactosamine (GalNAc) which are building blocks of larger polysaccharides. While GlcNAc and GalNAc commonly represent the terminating amino sugars of microbe-associated polysaccharides, such antigens are absent or exhibit limited serum accessibility within the human glycome.^6^

The recent application of printed glycan arrays to the study of serum antibodies has established an expanded view of the saccharides targeted by ACAs. Notably, these studies have revealed that Abs reactive with (GlcNAc), β-GlcNAc, or polymers of β1-4 GlcNAc such as found in chitin oligosaccharides are highly abundant in most individuals.^3,7,8^ These array-based approaches to dissect the global reactivity of serum ACAs have, in many ways, confirmed long-held views that were built by exhaustive investigations of individual carbohydrate antigens. Yet they cannot deconvolute the true heterogeneity of ACA composition and offer no insight into the development and maintenance of the B cells responsible for ACA production. For example, it is not clear if the same ACAs within an individual bind structurally similar chitins and monomeric GlcNAc, or if the ACAs binding to these structures represent discrete pools of Abs. Because ACAs target ubiquitous sugars ‘in context’, a better appreciation of the promiscuity or focusing of the ACA constituents toward these structures would help elucidate their functions and provide clues into the development and maturation of ACA repertoires. Although the origin of B cells responsible for nAbs to widely conserved antigens such as glycolipids and carbohydrates is still being refined, studies in mice clearly demonstrate canonical serum IgM antibodies reactive to phosphorylcholine and phosphatidylcholine are present in germfree mice.^9–11^ These and other similar findings have fueled a prevailing view that nAbs arise in the absence of exogenous antigen. However, most serum ACAs, which are present at low or undetectable levels in GF mice, are generated and maintained in response to commensal bacteria thereby defining ACAs as a class of nAb dependent on microbial exposure.^2,12,13^ These observations, coupled with the rapid kinetics of ACA emergence following birth suggest that microbial exposure is also responsible for ACA induction in humans.

Using direct antigen-labeling and comprehensive phenotypic characterization to closely track the development of carbohydrate-reactive human B cells in postnatal tonsillar tissues and pediatric peripheral blood, we show that circulating GlcNAc-reactive B cells emerge during early life and rapidly acquire memory B cell (BMem) phenotypes. Single-cell B cell receptor cloning and recombinant antibody expression coupled with profiling of gene expression in antigen-specific single B cells, show that GlcNAc-reactive B cells undergo GC-dependent clonal expansion, accompanied by BCR diversification evolving antibodies of demonstrable higher affinity. We also find that within an individual donor multiclonal heterogeneity is responsible for a distinctive antibody fine specificity signature for multiple discrete GlcNAc-containing structures. However the individual repertoires across multiple donors converge on a common distribution of anti-GlcNAc reactivity profiles. Moreover, similar reactivity profiles were encoded by private Ab gene rearrangements across individuals. Taken together our results provide a mechanism by which encounters with microbial antigens establish and diversify the systemic human ACA repertoire.

## Results

### B cell immunity directed toward N-Acetyl-Glucosamine emerges postnatally

To better resolve the emergence of humoral immunity to GlcNAc-bearing polysaccharide antigens in humans, we evaluated sero-reactivity to the GlcNAc-rich *Streptococcus pyogenes* (Lancefield Group A Streptococcus [GAS]) Carbohydrate (GAC) antigen in plasma isolated from umbilical cord blood, together with peripheral blood from juveniles, adolescents, and adults. All pediatric and adult plasma samples exhibited detectable IgM Abs against GAC, as well as class-switched antibodies of predominantly IgG, and to a lesser extent, IgA isotypes (**Figure 1a, S1a,**). Cord blood contained much lower levels of GAC-reactive antibody which was restricted to IgG, indicative of maternal origin as has been described for ACA-reactive IgG.^14^ Consistent with previous reports regarding GlcNAc sero-reactivity^15^, we observed that GAC-specific IgM titers were highest in adolescence and decreased moderately into adulthood (**Figure 1b**). We further observed strong correlations between IgM and class-switched IgA/IgG antibody titers within individuals, suggesting their concomitant development during immune exposures of early childhood (**Figure S1b-d**), as has been also observed with other ACA specificities.^4^

**Figure 1.**
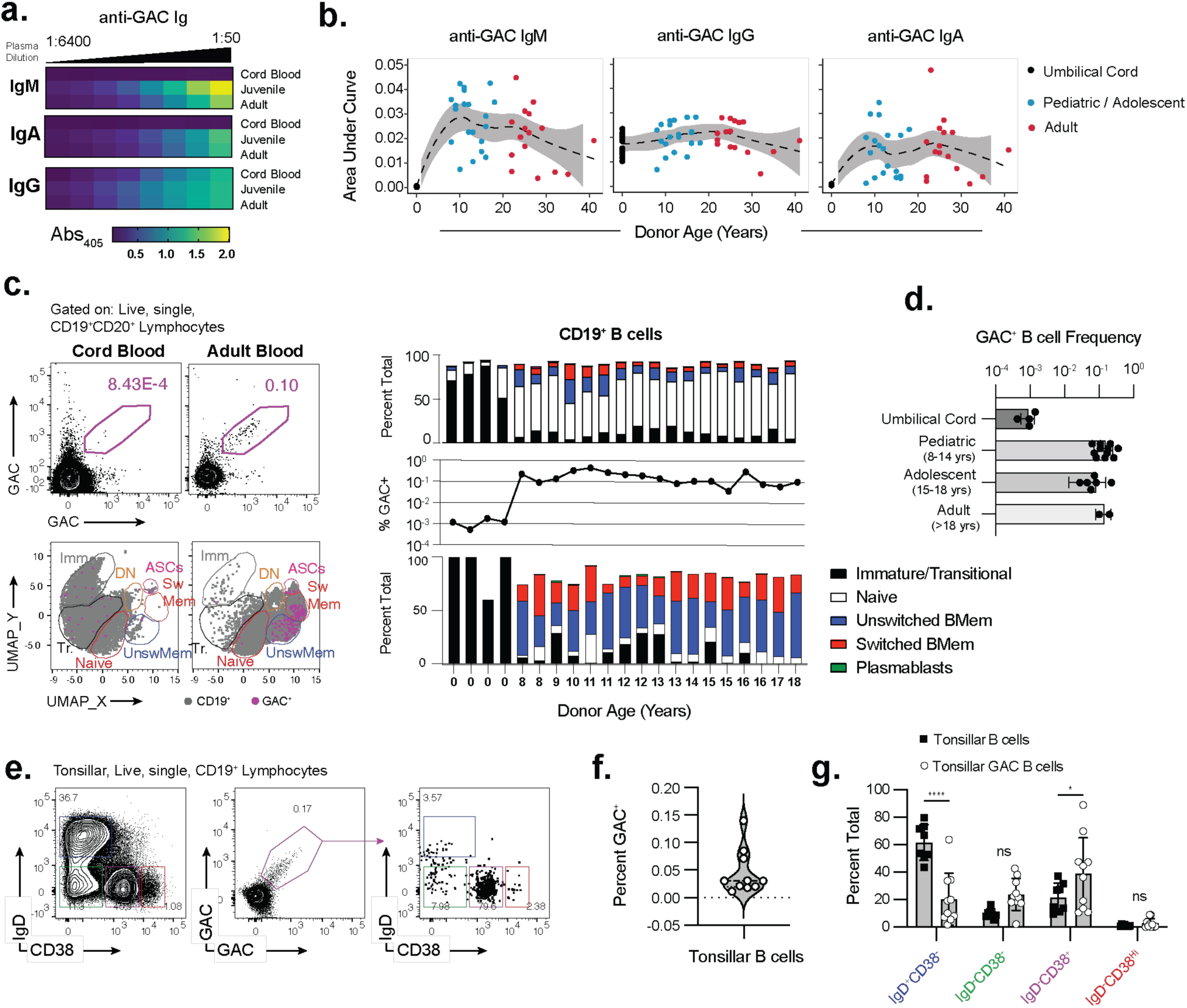
Emergence of GAC^+^ B cells and antibody in humans. **(a)** Heat map of ELISA absorbance values of anti-GAC antibody of the indicated isotype in cord blood or serum. Measurements were performed in triplicate for each serial dilution. **(b)** Area under the curve values of GAC^+^ Ab of the indicated isotype plotted by donor age. **(c)** GAC^+^ B cells visualized in cord or adult blood. GAC^+^ gating (left, upper) and UMAP projection of the 11 antibody panel to identify the labeled cell types with GAC^+^ cells overlaid (left, lower) (see methods). Representative flow histograms of GAC^+^ (left, upper) and subset distribution (left, lower) from the indicated source. Frequency of B cell subsets (right, upper) or GAC^+^ B cells (right, center and lower) by donor age. **(d)** GAC binding B cell frequency from donors segregated by age. **(e)** Representative flow histograms of B cell subsets (left) and GAC^+^ B cells (center) and B cells gated on GAC^+^ (right) in tonsil. **(f)** Frequency of GAC^+^ B cells in 10 tonsil specimens. **(g)** IgD and CD38 expression in unfractionated and GAC^+^ B cells in 10 tonsil specimens. Statistically significant differences by two way ANOVA in frequency are indicated (* p<.05, *** p<.0001).

Direct visualization of GAC^+^ B cells within PBMC samples by flow cytometry^12^ revealed that they comprised low frequencies (∼0.1%) of B cells in pediatric, adolescent, and adult volunteers (**Figure 1c,d**). Notably, we found that human GAC^+^ B cells predominantly express markers associated with canonical memory B cell (BMem) phenotypes, even in pediatric donors as young as 8 years of age. While the majority of circulating GAC^+^ BMem were unswitched IgM^+^ B cells, ∼20% expressed class-switched memory phenotypes that included both IgA and IgG in concordance with our serological findings (**Figure 1c**). B cells capable of binding to labeled GAC^+^ were scarce in cord blood, and the few that were resolved displayed immature and transitional B cell phenotypes, in striking distinction to those of pediatric origin (**Figure 1c,d, S1e,f**). Within the range of our cohort, donor age did not correlate with the frequency of circulating GAC^+^ B cells in peripheral blood, nor the proportion of GAC^+^ IgM or class switched BMem (**Figure 1c**).

Collectively, these data demonstrate that human GlcNAc-specific serum antibodies develop after birth, concomitant with the emergence of circulating antigen-specific B cells that express BMem phenotypes and diverse immunoglobulin isotypes. These data are in accord with previous reports describing the kinetics of GlcNAc-reactive Ab emergence in early adolescents^15^, and our correlation of their accumulation with the establishment of GlcNAc-reactive BMem cells highlights a key role for antigen experience in the establishment of this repertoire.

### Juvenile tonsillar GAC^+^ B cells express memory and germinal center phenotypes

Peripheral GAC^+^ B cells in the youngest pediatric samples we evaluated already exhibited adult-like BMem phenotypes which precluded our ability to make inferences regarding their development. Given that we previously established mucosal tissues as important drivers of GAC^+^ B cell development in mice^12^, we reasoned that tonsil tissues, which are considered an analog of Peyer’s patches for the upper aero-digestive tract and commonly encounter GlcNAc-bearing organisms including GAS, may harbor emerging GAC^+^ B cells.^12^ We therefore evaluated the frequency and cell surface phenotype of GAC^+^ B cells in n=10 tonsil specimens, collected from pediatric patients undergoing tonsillectomy at Children’s of Alabama through the UAB Tissue Biorepository. Despite the relatively young median age of our pediatric donors (median donor age 2.5 yrs, range 0-11 yrs), we consistently detected GAC^+^ B cells in these tonsil specimens. Intriguingly, a subset (3/12, 25%) of patients exhibited high frequencies of GAC^+^ B cells (7% to 15%) (**Figure 1e, f**). Again, donor age did not correlate with the frequency of GAC^+^ B cells. Although we were blinded to the clinical indication for each patient undergoing tonsillectomy, a portion of all pediatric tonsillectomies occurs due to Recurrent Tonsillitis, which is commonly associated with infection by GAS.^16^ Thus, it seemed likely that the expansion of GAC^+^ B cells observed in a subset of donor specimens was associated with ongoing or recent infection by GAS.

In contrast to circulating GAC^+^ B cells, which were homogeneously BMem, tonsillar GAC^+^ B cells exhibited greater phenotypic heterogeneity (**Figure 1g**), and included a minority of IgD^+^ naïve B cells as well as CD38^Hi^ ASCs. However, a large portion of GAC^+^ B cells displayed IgD^-^CD38^Hi^ phenotypes, consistent with that of germinal center (GC) B cells (**Figure 1e, g**). The striking correlation between the frequency of GAC^+^ B cells and the proportion of GAC^+^ GC B cells suggested that increased frequencies of GAC^+^ B cells directly resulted from GC expansion (**Figure 1g**). As the youngest donor we evaluated was less than 1 year old, these observations demonstrate that GlcNAc-reactive B cells are recruited into immune responses very early in childhood, and further implicate mucosal sites in the formation of B cell memory toward GlcNAc as a consequence of environmental microbial exposures.

### GAC binding B cells utilize receptors with discrete GlcNAc reactivities

Unlike mice, humans are frequently exposed to GAS. Although the GAS cell wall preparations we used to identify GlcNAc-reactive B cells in mouse systems exhibit high fidelity^12^, we sought to confirm that B cells labeled with GAC were reactive to, and specific for GlcNAc epitopes. We therefore performed single-cell sorting of GAC^+^ B cells from five of the characterized pediatric tonsil specimens including the three samples that exhibited expanded frequencies of GAC^+^ B cells. We then amplified, and cloned their respective Immunoglobulin Heavy Chain Variable (IGHV) and Immunoglobulin Kappa (IGKV) or Lambda (IGLV) light variable chains genes into mammalian expression vectors. We co-transfected pilot-scale (1mL) cultures to generate recombinant monoclonal Ab (rmAbs) and collected supernatants for specificity screening using a cytometric bioparticle/bead array. This array comprised of GAS and *Streptococcus dysgalactiae* (Group C Streptococcus [GCS]) which contain the native cell wall polysaccharides of these organisms, along with GlcNAc neoglycoconjugate coupled beads and anti-HuIg capture beads to visualize rmAb (**Figure 2a**).

**Figure 2.**
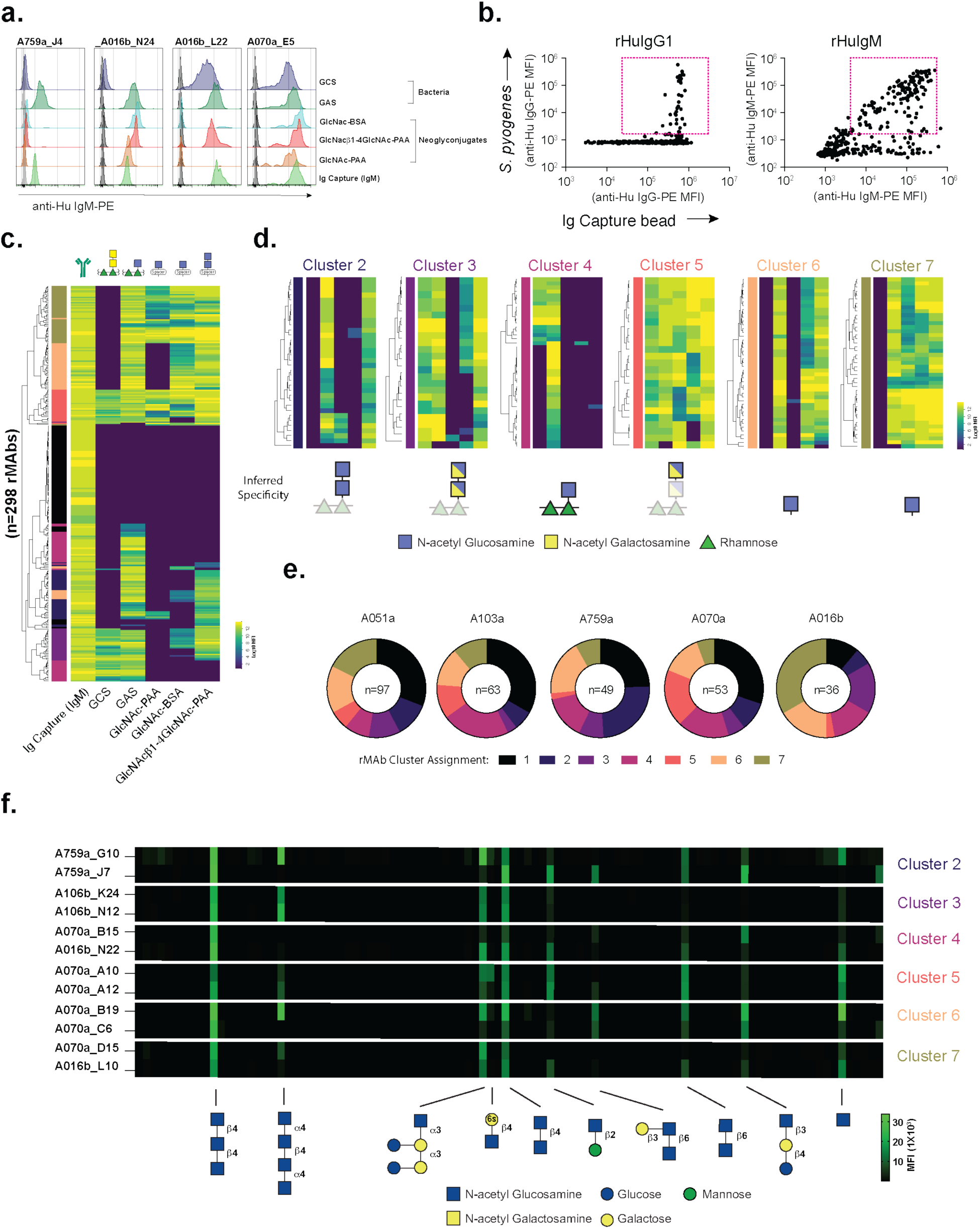
GAC^+^ B cell receptors display heterogenous binding to GlcNAc containing structures. **(a)** Representative flow cytometric histograms of four single-cell derived rhIgMAbs binding to the indicated elements of the cytometric array. **(b) g**MFIs of GAS stained with single-cell derived rmAbs expressed as IgG1 (left) or IgM (right). See **Figure S2**a for isotype comparisons and **Figure S2b** for polyvalency correlation. Red dashed boxes indicate positive GAS binding. **(c)** Heatmap of the gMFI of rhIgM derived from GAC^+^ tonsillar B cells on the indicated array elements. Hierarchical clustering dendrograms are shown on left side. Left color bars indicate Kmeans reactivity cluster designation (1-7). See **Figure S2c-d** for kmeans clustering plots and distance matrix. Schematics of antigen structures above. **(d)** Heatmap of the gMFI of rhIgM of the indicated reactivity clusters. Array element order is conserved from (c). Inferred glycan antigen schematic indicated beneath. **(e)** Distribution of reactivity clusters within rmAbs isolated from the indicated tonsil specimens. Number of rIgM recovered indicated in center of donut plots. Reactivity cluster designation indicated by color. Ages of donors: A051a <1yr, A103a 2yr, A759a 2yr, A070a 3yr, A016b 5yr. **(f)** Heatmap of the gMFI of members of reactivity groups 2-7 assayed on printed 100 glycan arrays. Measurements of the indicated rhIgG1_hex_ binding antigens printed in quadruplicate. The structure of bound glycans is indicated below.

Initially, we utilized human IgG1 Fc sequences to mediate recombinant expression of our GAC-reactive antibody constructs. However, despite providing high recovery rates of rmAbs, few exhibited detectable binding to GAS or GlcNAc-haptenated antigens, and detectable GAS-binding was notably limited to those clones with the highest rmAb expression levels in supernatant (exceeding 5ug/mL) (**Figure 2b**). We speculated that anti-GAC rmAbs were likely of low affinity, and perhaps below the limit of detection when expressed as a bivalent hIgG1. To determine whether increased valency improved our recovery of GlcNAc-binding antibodies, we generated a recombinant human IgM heavy chain expression construct which, when expressed in the absence of a J chain, result in the production of hexameric IgM molecules (rHuIgM).^17^ Indeed, this modification resulted in a four-fold increase in the frequency of rmAbs binding to GlcNAc-bearing antigens compared to rIgG1 (**Figure 2b**). We further confirmed that the difference in reactivity of these rmAbs is predominantly due to valency, as most of these antibodies also exhibited equivalent binding to GAS when expressed in a distinct hexameric IgG1 format^18^ (**Figure S2a, b**).

Analysis of rHuIgM binding profiles using the cytometric bioparticle/bead array revealed significant heterogeneity in their reactivity toward different configurations of GlcNAc. While some rmAbs bound only to GAS, others recognized GlcNAc haptens presented on both bovine serum albumin (BSA) and polyacrylic acid (PAA) backbones. Surprisingly, we identified antibodies that bound GCS, which bears a distinct cell wall carbohydrate decorated with terminal di-N-acetyl-Galactosamine^19^ residues in addition to binding GAS and GlcNAc-haptenated structures, suggestive of broad reactivity to N-acetylated hexosamine epitopes (**Figure 2a,c,d**). When we considered the reactivities of n=298 recovered Abs in aggregate, distinct reactivity profiles emerged. k-means clustering on the intensity of antigen binding of culture supernatants, measured by geometric mean intensity (gMFI) of anti-hIgM PE secondary, against each element of the array identified 7 distinct clusters which were segregated by dimensional reduction visualization and distance space (**Figure S2 c,d**). Additionally, these clusters were well resolved when the MFI tables were subjected to unsupervised hierarchical clustering (**Figure 2c**).

While the members of the largest cluster, cluster 6 (n=45) exhibited binding to monomeric GlcNAc, GlcNAc-BSA, chitobiose, and GAS, they did not bind GCS or GlcNAc-PAA. (**Figure 2d**). The smallest clusters (3 and 5), n=25 display binding to GAS as well as GCS, and in the case of cluster 5, every other form of GlcNAc represented in the array (**Figure 2d**). Remarkably, each of these reactivity clusters was represented in every patient specimen (**Figure 2e**). We next sought to confirm the broadly hexosamine-specific binding profiles observed using the cytometric array and evaluate binding to a greater breadth of carbohydrate targets. We therefore selected two representative Abs from each ‘specificity cluster’, to examine their global reactivity as hexameric IgG1 molecules on microarrays printed with 100 glycan antigens (**Figure 2f**). Of the twelve mAbs tested, all exhibited restricted reactivity to GlcNAc terminating glycans. Each Ab bound chitotriose (polymers of β1-4GlcNAc) while all except A759a_J7 exhibited strong binding to monomeric GlcNAc, thereby confirming the specificity detected by our cytometric array. This analysis also revealed significant differences in the fine specificities of these rmAbs, even between members of the same specificity cluster. For example, cluster 7 member A070a_D15 tolerated terminal α4 linkages in GlcNAc polymers whereas A016b_L10 did not. Conversely, A016b_L10 demonstrated reactivity toward β1-2 and β1-6 GlcNAc that was not observed in A070a_D15, despite equivalent recognition of chitobiose and chitotriose by the two clones (**Figure 2f**).

The expression of rAbs derived from GAC^+^ B cells as polymeric Ig rather than bivalent IgG1 greatly increased the sensitivity of binding assays, presumably by impacting avidity of the antibody-antigen interactions, and revealed substantial heterogeneity in GlcNAc-reactivity among patient-derived rmAbs. Similar to our previous study of GlcNAc-specific Abs in mice^20^, human anti-GAC antibodies demonstrate restricted and discrete reactivity towards GlcNAc-containing glycans but differ from one another in their reactivity patterns or relative intensity toward individual glycan targets. These data further highlight an unappreciated level of heterogeneity among GlcNAc-reactive Abs in humans.

### GlcNAc reactivity is encoded by donor-specific, diversified receptors

When the sequences of the IGH and IGK/L genes of these clones were examined, these antibodies were encoded by 27 IGHV subgroups including members of IGHV1,2,3,4 and 5 gene families (**Figure 3a**). The most dominant clones represented between 18.5% to 27% (**Figure S3**) of IGHV sequences from a given specimen. Clonotypes from each sample were unique to that individual (**Figure 3b**), and nearly every clone exhibited IGHV somatic mutations (99.6%) with replacement mutations exceeding silent mutations by >3 fold (2.79-3.5) (**Figure 3c**). Individual members of expanded clonotypes exhibited variation in the number of somatic mutations within their sequences, which were unevenly distributed across the individual domains of the IGHV genes (**Figure 3d**). Considering the frequent GC phenotype of clonal members, these observations suggested that they were actively undergoing SHM. To evaluate this possibility, we used germline-weighted protein parsimony algorithms, implemented in Phylip^21^, to generate phylogenetic trees of the IGHV of members of two large lineages within the sequences of cloned Abs: lineage 132 (lin132) from A051a and lin192 from A070a. In both cases, we had cloned IGHV genes that accumulated sequential mutations across the evolutionary trajectory of the clonotype. For lin132, we had further recovered clones that originated as IgM expressing cells and were anchored earlier in the phylogenetic trees than those originating as IgG, supporting the hypothesis these cells were actively undergoing SHM and recent class switch recombination (**Figure 3e**).

**Figure 3.**
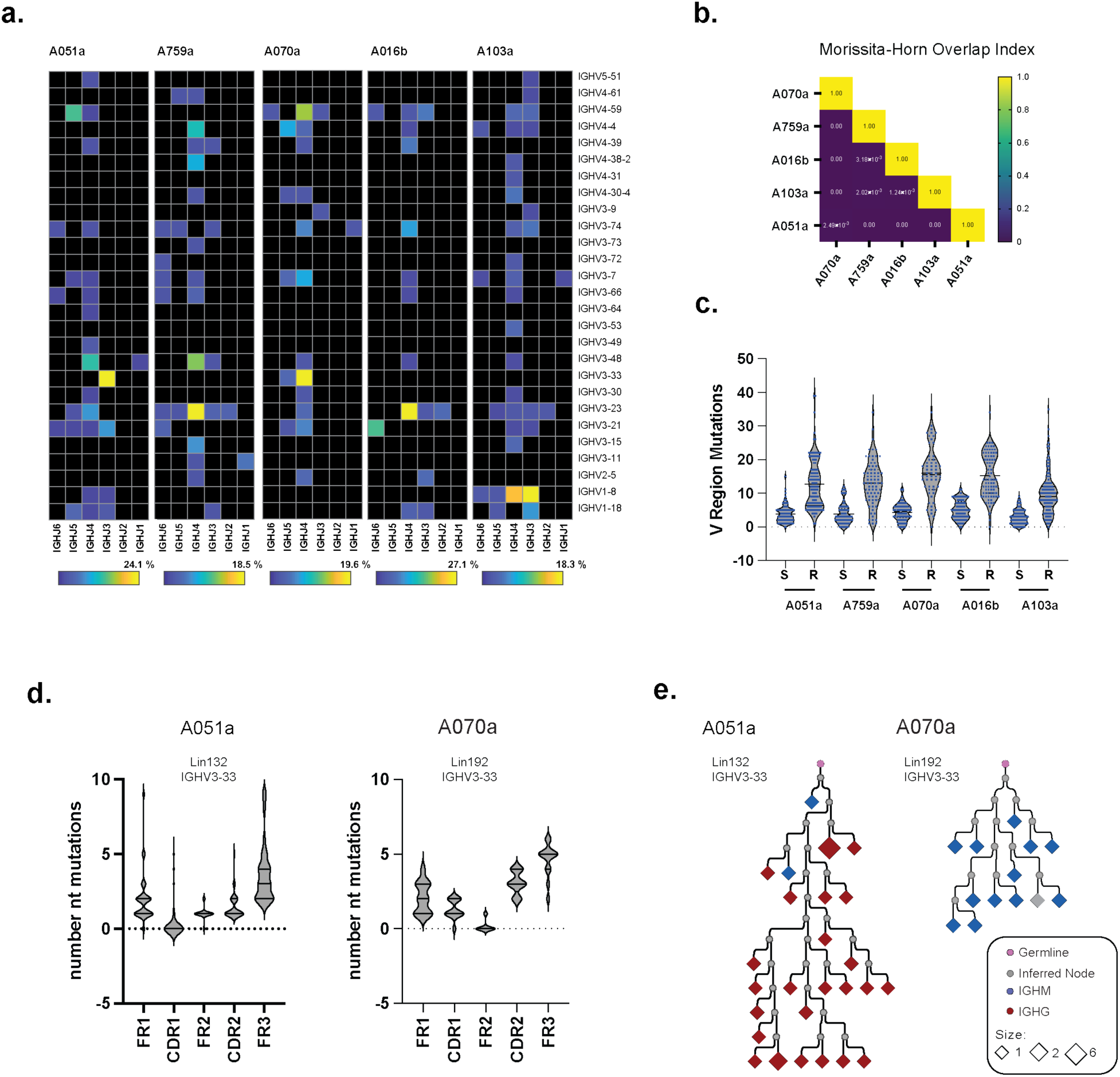
Tonsillar GAC^+^ B cell receptors are diverse, private, and somatically mutated. **(a)** Heatmap of frequency of IGHV:IGHJ rearrangements among GAC^+^ B cell receptors isolated from the indicated tonsil specimen. See **Figure S3** for specimen clonality plots. Color mapping represents percentage of recovered IGHV sequences per specimen. **(b)** Heatmap of inter-sample Morissita-Horn overlap distance matrix of IGHV clonotypes of GAC^+^ B cell receptors derived from the indicated specimens. **(c)** Violin plots with dot plot overlays of the number of silent (S) or replacement (R) nucleotide mutations of all IGHV sequences derived from the indicated tonsil specimens. **(d)** Violin plots of the number of nucleotide mutations in the IGHV domains of all members of the indicated expanded lineages. IGHV gene usage, specimen identifier, and lineage designation are shown. **(e)** Phylogenetic trees depicting the phylogentics of clonal members of the indicated expanded GAC^+^ lineages. Specimen identifier, lineage number, and IGHV usage are shown. Clones are colored by originating isotype as indicated in key.

Collectively, these results suggest that in addition to expressing the pre-requisite surface phenotypes, GAC^+^ B cell receptors exhibit the sequence features consistent with a GC-dependent evolutionary trajectory. That we observed private (donor-specific) BCR sequence repertoires within the GAC-binding B cell compartment, but shared distribution of fine-epitope antibody specificity profiles, suggest a convergence of human anti-GAC repertoires with similar reactivity profiles but derived from distinct founder/parental sequences.

### Tonsillar GAC^+^ B cells participate in GC

Because carbohydrate reactive B cells are not often considered to be associated with the T-cell dependent processes of the GC, we sought to evaluate the transcriptional profile of GAC^+^ B cells in tonsils as a means to validate their classification as GC B cells. We selected two patient specimens, <1yr (A051a) and 2 yrs (A103a), that exhibited an intermediate expansion of putative GAC^+^ GC B cells and sorted patient-matched CD19^+^ and GAC^+^ B cells for single-cell RNA-sequencing (scRNA-Seq) using the 10x genomics platform. We sequenced 5’ gene expression (5’ GEX) and V(D)J libraries for each cell population and performed dimensional reduction analysis by uniform manifold approximation and projection (UMAP) (**Figure 4a**).

**Figure 4.**
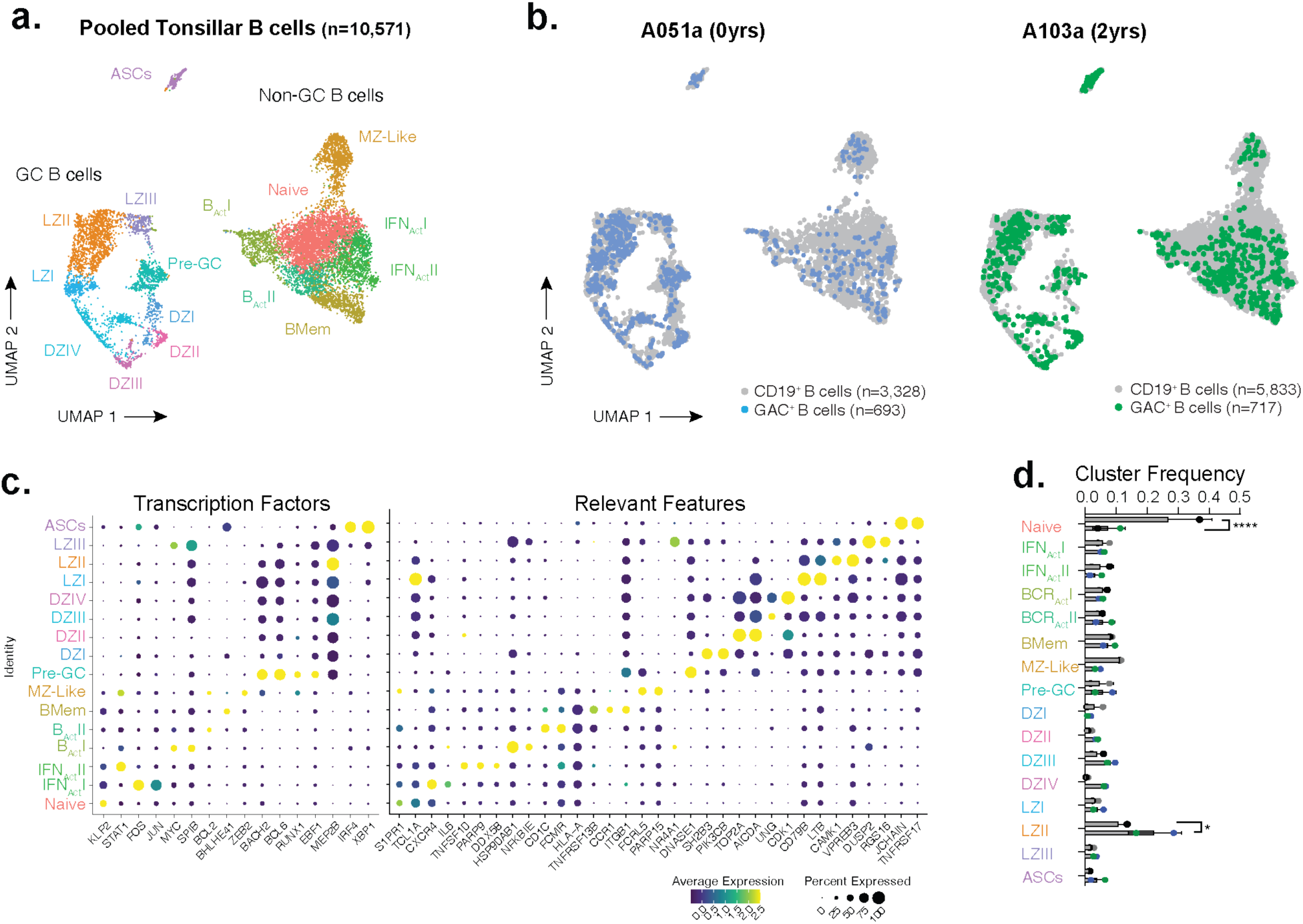
Tonsillar GAC^+^ B cells are distributed across subsets but enriched in germinal center B cells. **(a)** UMAP projections of integrated Seurat object including unfractionated and GAC^+^ B cells from specimen A051a and A103a. Transcriptionally defined cell clusters are indicated by color and labeled by their annotated subset identity. See **Figure S4a** for differentially expressed genes heatmap. **(b)** Overlay of UMAP projections of all B cells and GAC^+^ B cells of the indicated specimen. See **Figure S4b** for isolated GAC^+^ B cell UMAP. Cell numbers for each sample are shown. **(c)** Feature plot of the differentially expressed transcriptional regulators (left) and genes (right) that define B cell cluster identity annotation. See **Figure S4c** for feature plots of relevant gene expression in UMAP space. **(d)** Frequency of unfractionated or GAC^+^ B cells assigned to each B cell subset cluster. Bars represent means, dots are colored by source. Statistically significant differences by two way ANOVA in distribution between unfractionated and GAC^+^ B cells are indicated. (* p<.05, **** p<.0001).

In agreement with other reports^22,23^, this analysis identified many transcriptionally distinct B cell clusters across all four libraries comprising three non-contiguous superclusters that were broadly representative of germinal center B cells (GCB), non-germinal center B cells (non-GCB), and antibody-secreting cells (ASCs) (**Figure 4a**). The non-GCB population consisted of seven transcriptionally distinct populations we classified as naïve, MZ-like, Atypical BMem (AtypBMem), Activated B cells (B_ActI_ and B_ActI_), and BMem (**Figure 4b,c S4a,b**).The GCB supercluster further comprised eight distinct populations including Pre-GC B cells, four proliferative dark-zone (DZ) B cells clusters, and three clusters exhibited high expression of genes typically associated with light zone (LZ). (**Figure 4b, S4a-c**).

Each cluster was represented in both the CD19^+^ and GAC^+^ B cell library of each patient (**Figure 4b, d, S4b**). In agreement with our flow cytometry analysis, high proportions of GAC^+^ B cells were assigned to the GC supercluster, and were significantly enriched within the major LZ GCB cluster, while being highly under-represented in the naïve cluster as compared to bulk CD19^+^ B cells. Thus, these scRNA-seq data support the cell surface marker expression analysis confirming that GAC^+^ B cells acquire GC B cell phenotypes in tonsil tissues of pediatric patients.

### GAC^+^ B clonotypes are differentially expanded and diversified

We leveraged the Ig-Seq analysis pipeline to facilitate IMGT-based annotation of 10X-derived immunoglobulin heavy- and light-chain VDJ/VJ sequences to define clonal lineages of B cells.^24^ We obtained a total of 5,841 productive immunoglobulin heavy chain (IGH) sequence reads of varying immunoglobulin isotype distribution, representing 358-3,085 sequences per specimen (mean 1,460) (**Figure 5a**). The GAC^+^ B cells from both patients included multiple expanded lineages in which the dominant clonotypes represented 10% (A051a) or 20% (A103a) of all GAC^+^ B cell-derived IGH sequences (**Figure 5a,S5a**). By contrast, CD19^+^ B cell IGVH sequences consisted largely of singletons.

**Figure 5.**
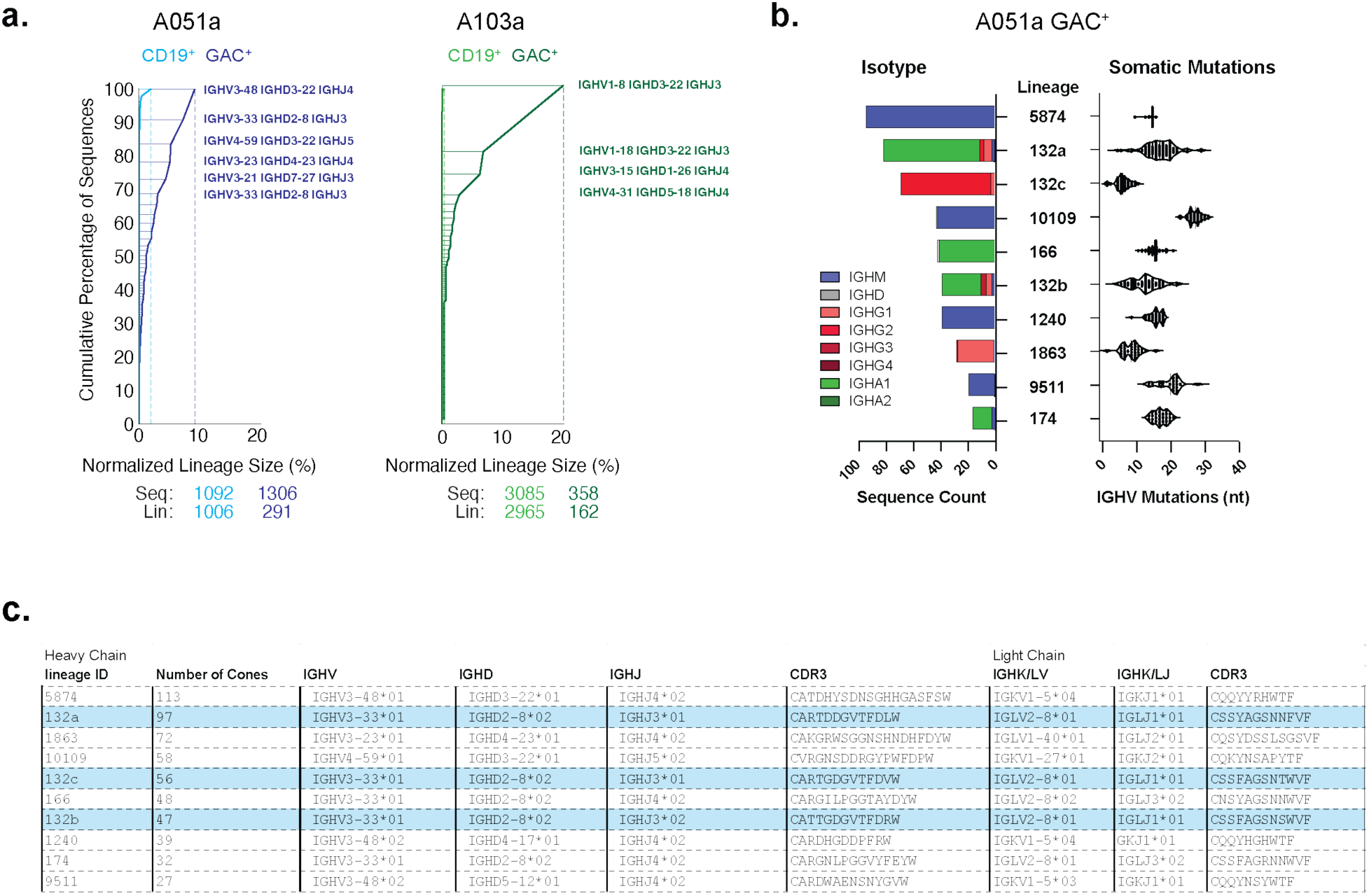
GAC^+^ B cell clonotypes are expanded and diversified in the tonsil. **(a)** Cumulative frequency of B cell clonal lineages expanded in the indicated specimens. Rearranged genes encoding most expanded GAC^+^ lineages are shown. See **Figure S5a** for heatmaps of IGHV:IGHJ frequency. **(b)** Isotype composition (left) and number of nucleotide mutations (right) of the ten most abundant lineages identified in A051a GAC^+^ B cell 10x sequences. See **Figure S5b** for unfractionated B cell and GAC^+^ B cell isotype frequencies. **(c)** Table of the indicated features of the ten most expanded clonal lineages in A051a. Highlighted lineages 132(a-c) are highly diversified resulting from the informatic assignment into three distinct members resolved in **Figure S5.** See **Figure S5c-f** for lineage 132 deconvolution.

Although most GAC^+^ B cells exhibited SHM, the degree of diversification within expanded lineages varied. Among the IGHV sequences derived from A051a, the largest clonotype, lineage 5874, consisting of 137 sequences, averaged ∼15 nt mutations per IGHV with a standard deviation of only 0.5 nt mutations. The mean substitution of the next largest lineage, lin132 clones was similar to that of lineage 5874 (∼17nt), however there was much higher variance, 5 to 29 nt mutations with a standard deviation of 4nt substitutions (**Figure 5b**). Interestingly, although GAC^+^ cells were distributed across IgM, IgA, and IgG isotypes (**Figure S5b**), most expanded GCB lineages were highly enriched for a single isotype (**Figure 5b**).

When the IGHV annotation of the ten largest clonotypes of each specimen was examined, we noted that A051a-derived sequences contained three lineages that shared IGHV and IGHJ gene segments, encoded highly similar CDR3’s and used similar light chains. In fact, when these sequences were aligned, we found they all shared 3 replacement mutations (a157c, t174a, g278t) which could not be explained as polymorphisms as they were only present in the IGHV3-30 of GAC^+^ B cells, and not the corresponding CD19^+^ B cells (**Figure 5c**). We reasoned that these lineages may have arose from a common ancestor that had undergone extensive diversification. To evaluate this possibility, we performed reclustering of all A051a derived IGHV3-30 sequences with junction lengths equivalent to that of lin132 (single-cell and 10x). This effort resolved four groups that largely correlated with isotype distribution and somatic mutation rates when the pairwise nt distance of the V region or CDR3 junction was calculated (**Figure S5c,d**). Interestingly, nearly every sequence derived from an IgG2 expressing clone contained a 2aa deletion in CDR1. To better understand the phylogenetic relationship of these cells, we performed DNA parsimony^21^ to construct phylogenetic trees and mapped lineage group assignments (**Figure S5e**) and originating isotype (**Figure 5f**) with CDR1 deletions. Groups 1 and 4 were well segregated. Group 1 almost exclusively contained the IgG2 expressing cells which carried the S32:Y33 deletion. Two of the other three groups were less well resolved. Despite this, three distinct branches were apparent in the phylogenetic reconstruction which deviated by early diversification events, as all of group 4 and a few members of group 3 carried a common S36R mutation **(Figure S5e,f)**. Based on this analysis, we assigned these clones into 132a-c lineages (**Figure S5f**).

These data demonstrate that within a single individual, multiple distinct clonotypes converge onto GlcNAc-reactivity which exhibit unique maturation trajectories. While the degree of clonal expansion of specific clonotypes does not directly correlate with the rates of SHM or isotype switching, all GAC^+^ B cell IGHV carry SHM characteristic of GC-dependent affinity maturation. In the case of lin132, multiple sublineages are apparent which clearly arose from a single common ancestor, which itself was mutated. Although it is not clear if the segregation of these sub-lineages is the result of their participation in anatomically discrete GCs or the recruitment of distinct precursors derived from previous immune responses, these sublineages clearly diversified independently. These data support the view that antigen driven immune responses shape the carbohydrate reactive B cell repertoire through the SHM-dependent diversification of precursors.

### Germinal centers support the clonal expansion of GAC^+^ B cells

To define the B cell subset identity of expanded B cell lineages in UMAP space, we integrated lineage assignments with the filtered Seurat object through the cell-specific barcodes. On average, we were able to associate immunoglobulin gene sequences with 47% of cells retained in the filtered Seurat object (range=27.2-79.5%) and performed integrated V(D)J and 5’GEX analysis for these cells (**Figure S6a**). In both patient specimens, large lineages of GAC^+^ B cells were highly enriched in the GCB and ASC superclusters compared to the non-GCB supercluster (**Figure 6a**). GAC^+^ lineages were enriched in the B_Act_ and BMem clusters, compared to the naïve, or AtypBMem clusters. Calculation of the Morisitta-Horn (MH) overlap index between patient specimens confirmed there was no clonal overlap between the GAC^+^ B cell repertoires of the two patients (**Figure S6b**), further supporting the single cell sorting observations. There was a high degree of conservation between the frequency of clonotypes isolated by single-cell receptor cloning and expression and the 10x datasets **(Figure S6c)**.

**Figure 6.**
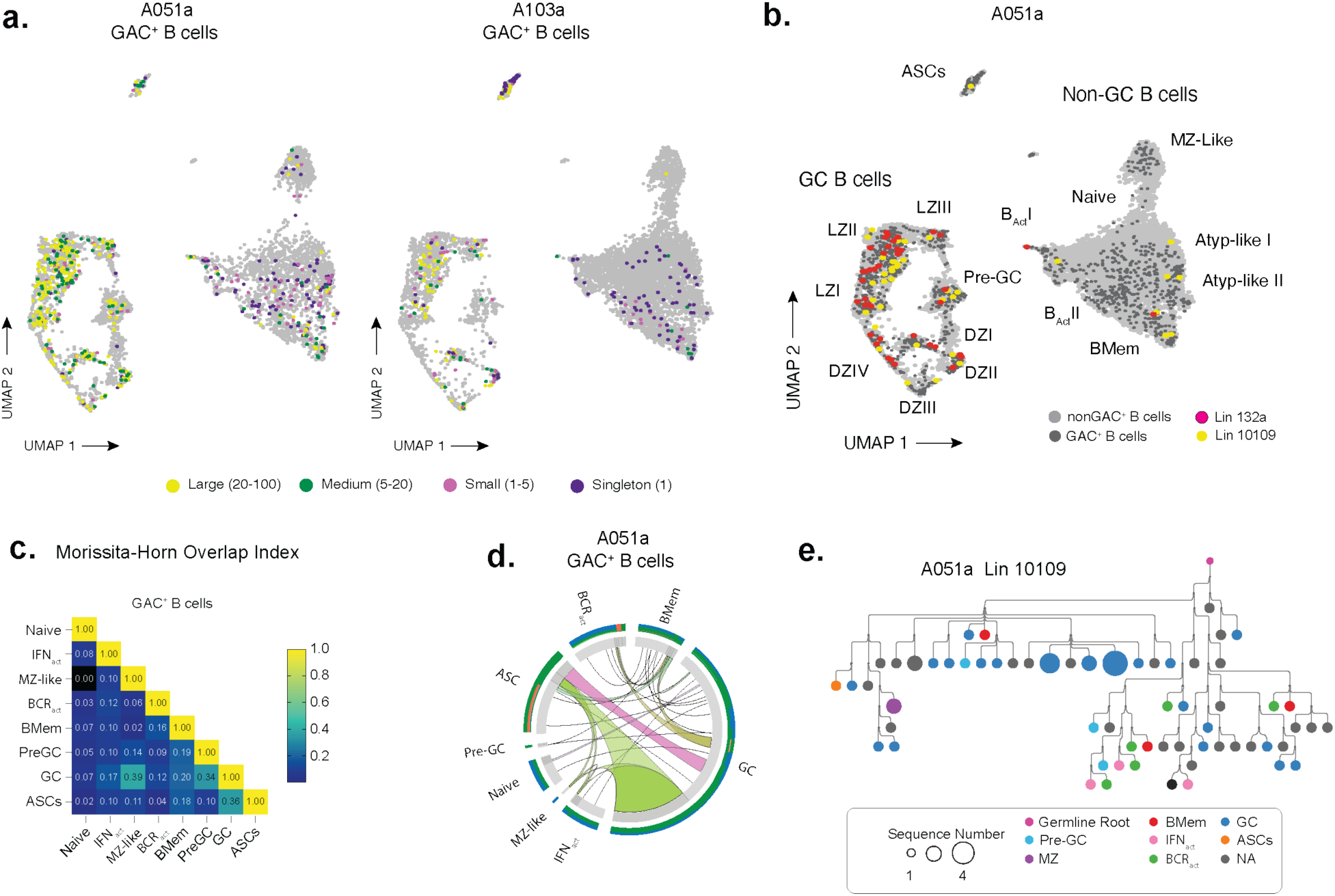
GAC^+^ B cells participate in GCs. **(a)** UMAP of expanded GAC^+^ B cells. All members of expanded lineages are colored by lineage size and overlayed onto unfractionated B cells in UMAP space. **(b)** Distribution of the indicated expanded lineages derived from A051a in UMAP space. **(c)** Heatmap of the Morissita-Horn overlap index of the GAC^+^ B cells assigned to the indicated B cell subset. **(d)** Ribbon diagram of GlcNAc-binding B cell clonal lineages identified in the indicated B cell compartments. Ribbons connect members of the same lineages across subsets. Colors assigned by cell subset connections. **(e)** Phylogenetic tree of lineage 10109. B cell subset identify denoted by color as indicated in key.

Some GAC^+^ lineages spanned the ASC and activated B cell clusters in addition to that of the GC B cells (**Figure 6b**). In fact, MH overlap measurements between cluster assignments suggested GAC^+^ lineages were highly shared across the Pre-GC GCB, and ASC B cell clusters within each patient (**Figure 6c**). Intriguingly, individual GC B lineages were variably connected to the BMem and MZ clusters, which were not themselves interconnected, suggesting MZ and BMem B cells maintained independent clonal identities following their GC participation (**Figure 6d**). Because some expanded clonotypes spanned multiple transcriptionally distinct B cell subsets, we again used phylogenetic trees constructed by protein parsimony to visualize the relationship between the diversified BCR sequence and cellular subsets within a single expanded lineage. We provide the analysis of lin1019, which included a large number of clones bearing transcriptomic profiles of varying cluster assignments. Despite most of the individual members of this lineage being categorized as GC B cells, BMem cells, activated B cells, and ASCs were distributed within the tree (**Figure 6e**), suggesting lin1019 MZ and BMem were progeny of GC B cells. Collectively these data demonstrate that GAC^+^ B cells are expanded in GC reactions and that both MZ and BMem GAC^+^ cells are progeny of expanded and diversified antigen-specific GC B cells.

### GAC^+^ GC B cells are affinity maturated

To determine if the somatic mutations observed in GAC^+^ B cell receptors altered the affinity and/or the fine specificity to GAC, we generated synthetic rhuIgM Abs derived from the consensus sequence of all members of the top 10 expanded lineages (**Figure 5a**) within the 10x A051a dataset as well as unmutated common ancestors (UCAs) of these lineages. We performed protein parsimony of all members of each lineage with the sequences of the consensus and UCA constructs and then generated phylogenetic trees (**Figure 7**). In the case of lin132, the consensus sequences fell within the phylogenetic reconstructions for each 132a, 132b and 132c sub lineage, while the UCA was closest to the non-rearranged IGHV3-33*01 gene. Except for lin1863, all consensus seq derived Abs exhibited strong binding to GAS. UCA derived Abs demonstrated significantly reduced binding to GAS compared to their corresponding consensus Abs (**Figure S7a**). When the MFI across a dilution of rhuIgM was compared, the MFI of UCA Abs at 5ug/mL was much less than that of the consensus sequences assayed a 40ng/mL for most lineages, indicative of a greater than 500 fold reduction in binding. The exception was lin132 UCA which was reduced 30-100 fold (**Figure S7a**). As with single-cell derived Abs, these lineage derived rhuIgM displayed differences in their reactivity to other hexosamine containing structures (**Figure S7b**). Because of the distribution of avidity observed between these rhuIgMs we sought to measure the kinetics of their association with GAC. Using GAS PGPS as a ligand, we subjected each purified rhuIgM to surface plasmon resonance analysis. Consistent with the flow cytometric analysis, consensus Abs and the UCA of lin132 demonstrated measurable association with GAC. However, the apparent K_D_ (K_Dapp_) of lin132c (1.9×10^-9^) was 10 fold lower than that of lin132a (2.9×10^-10^) and lin132 b (6.8×10^-10^). Lin132 UCA, which exhibited the highest binding to GAS of all lineage UCA’s by flow cytometry, was the only UCA that exhibited binding above the limit of detection at 8.3×10^-8^M, but exhibited a K_Dapp_ (5.7×10^-7^) that was 1000 fold lower that lin132a (**Figure 7b,c**). The other lineage consensus derived rhuIgM Ab’s exhibited K_Dapp_ values ranging from low nanomolar (9.7×10^-9^) to micromolar (4.0×10^-5^) (**Figure 7d,e**). Interestingly, the most expanded lineage, lin 5874, which was restricted to IgM, had the lowest binding affinity (**Figure 7d**). In every case, however, the consensus sequence rhuIgM displayed several orders of magnitude higher affinity than the UCA (**Figure 7f**) directly correlating with flow cytometric GAS staining by these rhuIgM’s (**Figure S7c**).

**Figure 7.**
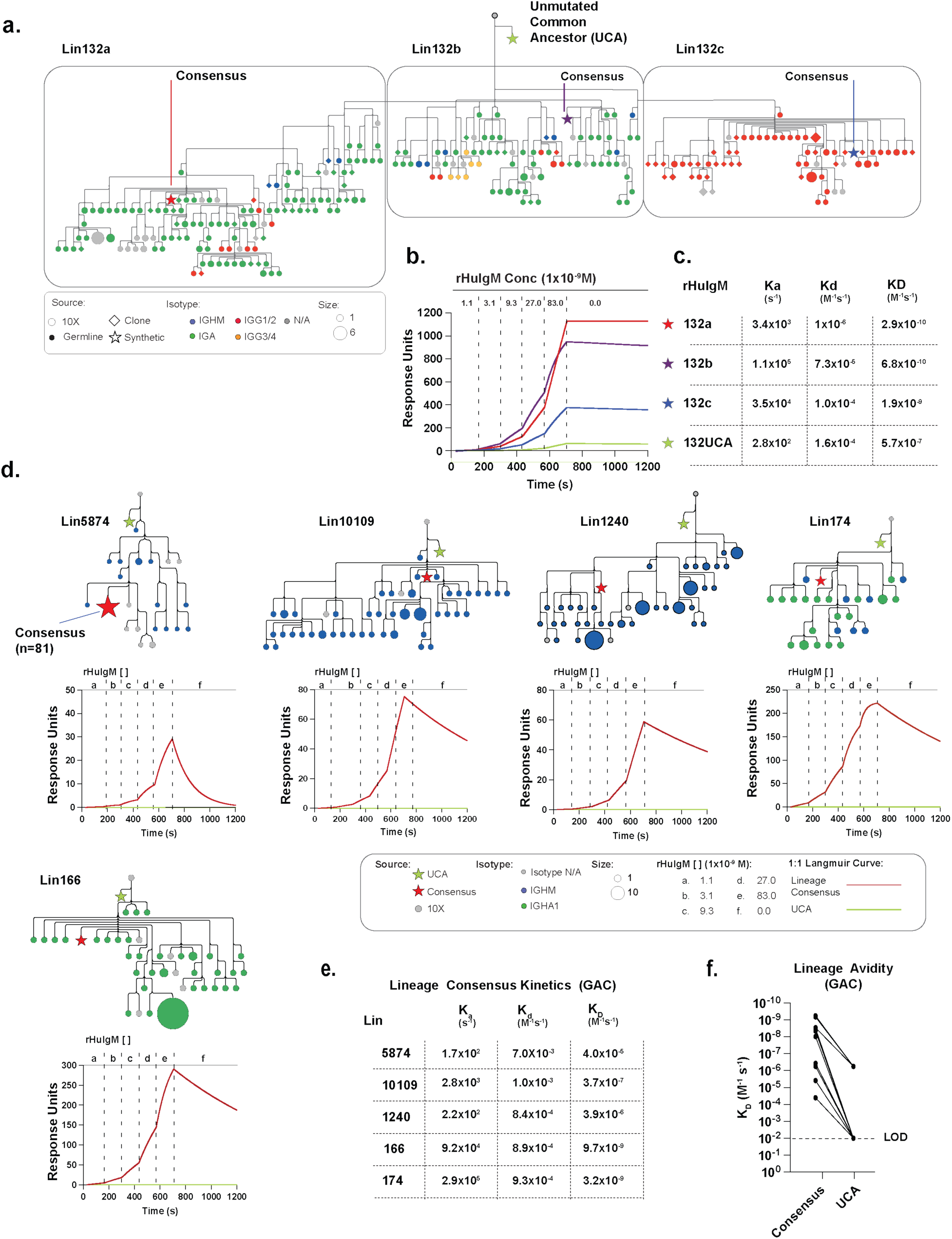
GAC^+^ B cells are affinity maturated. **(a).** Phylogenetic tree of all members of lineage 132 (a-c) with the positions of the consensus and unmutated common ancestor (UCA) indicated by stars. Individual clones are colored by isotype. See **Figure S7a-b** for consensus and UCA binding by cytometric array. **(b).** Spline plots of the 1:1 langmire curves from SPR at the indicated concentration of lineage 132(a-c) consensus and UCA rhIgM against PGPS. **(c)** Table of SPR derived association and disassociation constants and apparent affinity (K_Dapp_) of lineage 132(a-c) consensus and UCA rhIgM binding PGPS. **(d)** Phylogenetic trees of expanded GAC^+^ B cell receptor lineages from A051a. Position of consensus (red stars) and UCAs (green stars) are shown above spline plots for rhIgM binding to PGPS. Red traces are consensus sequences, green traces are UCA rhIgM Abs. Concentrations are indicated. **(e)** Table of SPR derived association and disassociation constants and apparent affinity (K_Dapp_) of the indicated lineage consensus rhIgM binding to PGPS. **(f)** K_Dapp_ of expanded A051a expanded lineage consensus and UCA rhIgM. See **Figure S7c** for correlation between binding assayed by cytometric array and SPR.

Collectively, these data demonstrate that the somatically mutated GC B cell receptors have significantly higher affinity than their ancestral, germline antigen receptors. When considered in the context of their clear clonal expansion, along with their GC B cell-like surface phenotype and gene expression signatures, these observations strongly suggest that GAC^+^ B cells gain affinity for carbohydrate antigens in the context of the GC.

## Discussion

Our findings demonstrate a clear role for GCs in supporting the proliferation and somatic hypermutation of carbohydrate-reactive human B cells. Although typical T-independent model antigens (NP-Ficoll and Dextran) initiate early phases of GC formation in mouse systems, they fail to sustain GC reactions, drive significant SHM, or generate high affinity Abs.^25,26^ Our data show that pediatric tonsil specimens contain a subset of GAC^+^ B cells transcriptionally indistinguishable from bulk CD19^+^ GC B cells, characterized by the expression of multiple genes associated with antigen-presentation and T lymphocyte co-stimulation. These observations stand in contrast to the prevailing view of carbohydrate-specific antibodies, stemming from mouse studies in which model T-independent antigen systems fail to faithfully recapitulate immune system activation as it occurs in response to antigenically complex microorganisms, which involve a multitude of covalently linked polysaccharides, phospholipids, and protein epitopes. In mice, GlcNAc represents a canonical nAb specificity. Although immunization with GAS will produce increased levels of anti-GlcNAc Abs.^12,20^ these Abs exist naturally in adult animals in the absence of vaccination. Rather the commensal microbiota is critical for both the production of serum Abs and the formation of the GlcNAc-reactive B cells which comprise this repertoire.^12^ Enhanced levels of GlcNAc reactive IgM resulting from GAS vaccination reduces the incidence of airway hypersensitivity to GlcNAc containing allergens and autoimmune T1 diabetes respectively.^20,27^ In both cases, GlcNAc-reactive Abs play a role in these diseases in part by promoting non-inflammatory antigen clearance mechanisms thereby limiting the effects of immune activation toward environmental and self-antigens: a hallmark of nAb function.

Glycan reactivities have been associated with innate-like B cells and mucosal responses in mice and humans since their discovery. In mice, GlcNAc-reactive B cells are phenotypically B1 B cells^12^, however whether humans possess an analogue of mouse B-1 B cells remains controversial.^28–31^ Carbohydrate reactivity in humans is typically associated with MZ-like B cells, which consist of unswitched BMem that exhibit important differences from their mouse counterparts. Unlike mouse MZB cells, human MZ cells include multiple subtypes and recirculate, as they are found in peripheral blood, tonsils, lymph nodes, and gut associated lymphoid tissue (GALT) in addition to the spleen. Also unlike mouse MZ B cells, human MZ B cells express antigen receptors that contain SHM.^32–34^ In agreement with these studies, GAC^+^ B cells are constituents of circulating switched and non-switched memory B cell compartments. We further observe GAC^+^ clonotypes distributed in MZ, GC, and BMem compartments *in situ*. Although many clonotypes are found in both the GC and MZ or within the Bmem and GC compartments, essentially no overlap was observed between the MZ and BMem clonotypes in either GlcNAc or CD19 cells. These data corroborate larger repertoire studies which have suggested that MZ and BMem cells converge in GC reactions while maintaining their cellular identity upon GC exit.^34^

How and why human innate B cell populations acquire SHM has been a long-standing question. There are multiple proposed models to explain how TI antigens, specifically those associated with commensal microbes may drive apparent affinity maturation. In the context of mucosal immunity, antigen-independent diversification of B cell receptors, coupled with the successive recruitment of higher affinity clones into TI immune responses in the GALT has been proposed as a mechanism to achieve affinity maturation in the absence of cognate T cell help.^35^ B cell prediversification, the antigen-independent alteration of BCRs through SHM or gene conversion is a mechanism of B cell repertoire formation in several characterized mammalian systems referred to as GALT species including rabbits and sheep. Multiple lines of evidence support that human B cell receptors undergo prediversification in the GALT.^36^ In this view, UCA of B cell receptors may not share specificity with their progeny, as antigen reactivity is a product not a driver of SHM. All GAC^+^ B cell lineages described here carry common mutations shared across all members of the lineage as shown for lin132. We observed a significant enhancement of K_Dapp_ by both somatic mutations and the expression of polyvalent Abs. Given these observations, the specificity of UCAs of carbohydrate-reactive Abs expressed as IgG1 should be interpreted with caution, as the valency of cell-surface expressed Ig complexing with highly repetitive carbohydrate structures, such as bacteria capsids or cell wall polysaccharides, is likely sufficient to drive B cell activation through antigen receptors with affinities too low to be measured in commonly employed immunoassays. Our study supports a hapten model whereby GlcNAc associated glycans are linked to microbial protein components which facilitate T cell activation when presented by GlcNAc-reactive B cells. Thus allowing glycan-reactive B cells to participate in GCs, where they undergo somatic mutation and affinity maturation in an identical fashion to TD antigen-reactive B cells. It is important to note, however, that while GAC^+^ B cells clearly participate in GC reactions, the multiple models to explain the accumulation of SHM are not mutually exclusive and may be employed in concert to generate a productive ACA repertoire.

Glycan reactive Abs are often described as exhibiting lower affinity than anti-protein Abs^37^. Indeed, many of the Abs described here fall below the limit of detection of flow cytometric assays as bivalent IgG1 even against the highly polyvalent GAC. The GC participation and significant diversification of GAC^+^ B cells raises the question of why higher affinities are not achieved. When considering the antigen, however, it is likely that an affinity ceiling exists for each clonotype. There are few structures of glycan reactive Abs complexed with their cognate antigen, and even fewer for anti-GlcNAc Abs.^37,38^ Considering the small size of GlcNAc (∼360d) and the limited number of residues associated with strong ionic bounds, the upper limit of affinity may be low. However, when expressed as polymeric constructs, in this case as hexameric IgM, the increased avidity provides K_Dapp_ equivalent to that of protein-reactive IgG.

Most described mAb’s reactive with GlcNAc were derived from mice immunized with GAS. A few however, including CDT110.6 (mIgM) and RL2 (mIgG1), which are commercially available reagents to visualize O-linked GlcNAcylation, were raised by immunization with a synthetic O-GlcNAc peptide^39^ and rat liver nuclear envelopes^40^ supporting the hypothesis that GlcNAc specificity extends beyond pathogen recognition. Consistent with the reactivity of GlcNAc rhuIgM cloned from tonsillar B cells described here, these antibodies display different reactivity toward GlcNAc containing structures. CTD110.6, as an example, binds most ²1-4 GlcNAc terminating structures, but it does not bind GAS.^20^ How GlcNAc-reactive Abs discriminate between highly similar structures is not clear. Given the diversity of Abs, there are likely many modes of interface between glycans and Abs as they clearly engage the small GlcNAc moiety expressed in different molecular conformations. Some Abs tolerate α-linkages, while others do not; some bind GalNAc, while most do not. An individual’s glycan-reactive repertoire is comprised of these individual Abs that have overlapping but discrete specificities which, along with the isotype distribution of these clonotypes, define the consequences of their interactions with the constellation of microorganisms and autoantigens incorporating GlcNAc motifs.

Like GALT, the palatine tonsil evolved to mediate immunity at a mucosal barrier site. Unlike other secondary lymphoid organs, antigen is transported directly into the lymphoid follicle by specialized antigen capture and transport mechanisms.^41^ And unlike lymph nodes, these barrier sites are characterized by continuous antigen loading and chronically active GCs. ACAs like the anti-GlcNAc Abs described here display a high degree of specificity for glycans often associated with microbes. Because carbohydrate antigens, including GlcNAc motifs exist on many organisms, B cells reactive to these structures may be iteratively recruited in immune reactions targeting a diverse array of antigens. Thus, the convergence of fine specificities among the GlcNAc-reactive Abs described here, may be the product of affinity maturation of diverse and private clonotypes toward the most frequently encountered but diverse epitopes. Perhaps the high frequency of Abs that are promiscuous for hexosamines, those reactive with GAS, GCS, and GlcNAc-neoepitopes were expanded in these individuals because of their apparent promiscuity. Thus, the conserved ‘universal architecture’ of the human ACAs may be derived from a lifetime of exposures and the selection for Abs that target the commonalities among rather than the uniqueness between microbial antigens.

### Limitations of this study

There are several limitations to this study. While we probed GAC^+^ B cells from five pediatric tonsil specimens, we performed single cell transcriptomics on only two. However, considering the degree of overlap between the single cell transcriptomic derived BCRs with those cloned by single cell sorting and the concordance of transcriptomic and cell surface phenotypes, we feel these numbers of samples and antibodies analyzed here is sufficient to draw generalizations about the human GlcNAc-reactive repertoire. It is also important to note, that all samples were derived from cryopreserved specimens, which may have led to uneven B cell subset recovery. Regardless, the transcriptomic profile of these specimens is consistent with previous studies.^22^ All tonsil specimens were derived from pediatric patients. Although we did not observe convergence of public B cell clonotypes, this aspect of the study may not accurately reflect the adult repertoire in which the GlcNAc-reactive clones may be further refined through a lifetime of exposure to microorganisms responsible for shaping these clones. Likewise, we did not examine any tissue other than tonsil. It is not known how the clonotypes described here relate to the systemic repertoires of the donors. In mice, there is significant overlap as our previous studies have specifically examined the relationship between the GlcNAc mucosal B cell responses and the systemic GlcNAc-reactive repertoire.^12^ Although we were blinded to the clinical indications for tonsillectomy for the specimens used in this study, it is possible, perhaps probable, that some of these patients suffered from recurrent tonsillitis from infections by GAS. GAS is a human pathogen whose infection may induce a heightened inflammatory state not reflective of homeostatic mucosal immunity. We did not conduct studies permitting a spatial segregation of GCs. Although there are clear immune phylogenetic relationships within GlcNAc-reactive B cell clonotypes described here, we cannot know if these cells were participants in the same GC reactions. In the case of lin132 for example, sub lineages may have been independently diversifying in discrete and segregated anatomical sites.

## Acknowledgments

We would like to thank Vidyasagar Hanumanthu, director of the UAB Flow Cytometry and Single Cell Core Facility, and it’s staff for their assistance with these studies. Additionally, we would like to thank Shanrun Liu, who operates the single-cell platform. We would like to thank Mike Crowley and the staff of the UAB Hefllin Genomics Core.

## AUTHOR CONTRIBUTIONS

R.G.K, J.F.K, and J.S.N. conceived the study. R.G.K and J.F.K. secured funding, oversaw project administration, participated in the original drafting of the manuscript and its revisions, and provided direct supervision. J.S.N, R.G.K conducted experiments, performed formal analyses, curated data, visualized data, wrote the original draft of the manuscript and participated in its revisions. A.F.R provided methodologic expertise and supervision. J.N.B., A.R.C., W.L.D, conducted experiments. R.G.K. and J.F.K supervised and facilitated project administration. R.S.D provided critical reagents. All authors reviewed and approved the final version of the manuscript.

## Materials and Methods

### Patient samples

Adult plasma samples were isolated from volunteers within the UAB research community under the IRB IRB-300002600. Pediatric plasma samples were provided by the UAB Endocrinology Research Clinic under the UAB IRB-300000100. Cord blood samples were collected form UAB woman’s and Children’s research clinic under the UAB IRB-170224007. Blood mononuclear cells were harvested by density gradient centrifugation with Histopaque following manufacture’s recommended protocols. Following centrifugation, plasma was harvest by pipetting and frozen at −80C until use. Mononuclear cells were isolated and washed with ice-cold PBS then cryopreserved in cell freezing media (10% DMSO, 50% FBS, in PBS). Surgical tonsil remnants were obtained from UAB Surgery tissue procurement. Tonsil specimens were mechanically dissociated using forceps, and passaged through 100um cell strainers (corning). Mononuclear cells were isolated as described above and cryopreserved until use.

### Enzyme-linked immunosorbent assays (ELISA)

The measurement of antigen-specific antibodies in donor plasma samples were performed as previously described in a 384-well ELISA format. Briefly, antigen stocks, [GAS 100S PGPS, Lee labs, GlcNAc-BSA PC-BSA (Vector labs) were diluted to working concentrations of 2 μg/ml and used to coat ELISA High-binding plates at 4°C overnight. The following day, unbound antigens were washed away and the plates were blocked with 2% Bovine serum albumin (BSA) at 37C for 1hr. Plasma specimens were diluted 1:50 in 96 well plates, serially diluted and arrayed into assay plates using a Biomek FX3. All samples were analyzed in triplicate. Following incubation, antigen-bound antibodies of the IgM, IgG and IgA classes were detected using the anti-human isotype specific alkaline phosphatase (AP)-conjugated goat anti-human secondary antibodies (Southern Biotech). Following extensive washing, bound antibody was by visualized through the addition 50ul of AP substrate (1mg/ml p-Nitrophenyl Phosphate (PNPP), in AP buffer (1 M diethanolamine pH 9.8)). Following the incubation, the reactions was stopped by the addition of an equal volume of 10N NaOH and absorbance of each reaction read at 405 on an OmegaStar plate reader. Area under the curve (AUC) analysis was used to quantitate the relative abundance of antigen-specific serum antibodies of each isotype across patients and to generate databases populated with patient metadata for comparative analyses between groups and longitudinal analysis across the cohort.

### Flow Cytometry and FACS sorting

Cryopreserved mononuclear cells isolated from peripheral blood or tonsil were rapidly thawed in a 37C water bath and non-viable cells were removed by density centrifugation over Histopaque. The cells were resuspended in ice-cold staining buffer (PBS, 1%FBS) and counted. The cell suspensions were then stained with antibodies: CD5, CD10,CD19,CD20,CD24,CD27, CD38, IgM. IgD, IgG,PGPS, FcRL5, CD83, CXCR4. For sorting, PGPS binding B cells were sorted on a FACSAria (BD Biosciences) in the UAB Comprehensive Flow Cytometry Core as single cells for recombinant monoclonal expression or in bulk populations for 10x transcriptomics. Data were analyzed in FlowJo v10.8.1 (BD Biosciences). For blood B cell phenotypic analysis, the following gating strategies were used: BMem CD19^+^CD20^+^CD27^+^CD24^+^CD38^-^, immature B Cell CD19^+^CD20^+^CD10^+^, transitional B cell CD19^+^CD20^+^IgD^+^CD27^-^CD24^+^CD38^+^.

### Generation of Bioparticle/Beads array

Custom cytometric arrays were generated as previously described^42^ using 4 µm and 5 µm carboxy functionalized array kits (Spherotech, PAK-4067-8K and PAK-5067-10K). Streptavidin (SA, Southern Biotech) was buffer-exchanged into PBS using the manufacturer recommended centrifugation protocol using PD-10 columns (Cytiva). Following exchange, SA was diluted to 2 mg/mL in PBS. SA was then conjugated to the beads by harvesting 5×10^7^ beads by centrifugation (7500 RPM for 5 min) and resuspended in 0.5 mL of the 2mg/mL SA in PBS. 0.5 mL of 6 mM 1-Ethyl-3-(3-dimethylaminopropyl)carbodiimide (EDC) in 0.05 M MES buffer pH 5.0 (Pierce) was then added and the reaction mixture was rotated at room temperature overnight. Following coupling, the reaction was quenched with 0.1 mL of 1 M tris pH 8.0 and incubated for 1hr at RT. The beads were then washed twice in 1 mL PBS and resuspended in PBS with 1% BSA and 0.005% NaN_3_ and stored at 4°C until antigen loading. To conjugate biotinylated antigens (GlcNAc-PAAb, Chitobiose-PAAb [GlycoNZ], anti-hIgM, anti-hIgG [Southern Biotech]), 1mg/mL solutions of each were incubated with SA-conjugated beads while rotating overnight at 4C. GlcNAc-BSA was directly conjugated to beads as described above. Heat killed, pepsin treated GAS J17A4 (ATCC 12385)and formaldehyde fixed GCS C74 (ATCC 12388) were prepared as described^43^, diluted to 1e8 particles per mL and labeled with Alexa-647 and Alexa-488 (Thermo Fisher) respectively.

### Recombinant monoclonal antibody cloning, expression, and sequencing

Recombinant monoclonal Abs (rmAbs) were generated as previously described.^44^ Briefly, cDNA was generated from mRNA isolated from single PGPS-binding B cells-sorted into hypotonic lysis buffer in 384-well plates which were stored at −80C unit processing. PCR IgV_H_ and IgV_K/L_ amplicons, generated from the single cell cDNA, were cloned into expression vectors containing the constant regions of human IgG1, hexameric IgG^18^, IgM, for IGHVs or IgK, or IgL for light chain variable regions. Plasmids were sequenced on the Oxford Nanopore Minion platform using indexing array barcodes. IgH and IgL plasmids were co-transfected using polyethyleneimine^45^ into 293FreeStyle cells in Freestyle expression media (Invitrogen). Supernatants were screened for Ig and antigen reactivity using the cytometric array described. Hexameric IgG was purified using Sepharose G Fast Flow (Cytiva) and rIgM was purified using IgM CH1 Capture select resin (Thermo Scientific). rmAb concentrations were determined by UV spectrophotometry (Nanodrop, Thermo Scientific).

### Cytometric Bead Arrays

Ab binding determinations by cytometric array were performed as previously described. Sample dilutions or culture supernatants were arrayed into 96 well round bottom plates (Costar). Beads/bioparticles were mixed to a concentration of 6e5 particles per mL and 5ul was added to each well. Following a 10min incubation at RT, 200ul of PBS was added to each well, and the beads were then isolated by centrifugation. Supernatant was removed by flicking, and the beads were resuspended in 25ul of the appropriate anti-Ig secondary antibodies diluted 1/400 in 1% BSA PBS. Following a 10 min incubation at rt, the beads were again washed in PBS, resuspended in 80ul of PBS and subjected to flow cytometric on a Cytoflex flow cytometer (Beckman Coulter Life Sciences) in plate mode. Data were analyzed in FlowJo v10.8.1 (BD Biosciences) and geometric mean fluorescence intensities (gMFI) of binding by each rmAb to each particle were determined.

### Glycan arrays

Analysis of rIgG1_hex_ binding to glycans was performed using the Glycan Array 100 (RayBiotech). The manufacturer’s recommended protocol was followed with minor modifications to accommodate IgG analysis. Briefly, slides were stained with 2ug/mL rmAb in supplied buffer for 2hrs with gentle rocking. Slides were washed and stained with Alexa-488 labeled anti-human IgG (Southern Biotech) at rt using supplied buffers. Following a 2 hr incubation, the slides were sequentially washed with wash buffer followed by distilled water. Slides were air dried at rt in the dark overnight. Ab binding was visualized using a GenePix 4000B imager. Average MFI of the quadruplicate printed glycans were calculated in Prism (graphpad).

### Bioinformatics and phylogenetic analyses of IgV_H_-Seq data

IgV_H_ sequencing data were processed and analyzed as described previously.^24^ For 10x derived data, paired-end reads were assembled and quality filtered using FastQC scores. Compiled Ig sequence fasta files were submitted to IMGT/HighV-QUEST^46^ for annotation. Annotated sequences were filtered for productivity prior to further analysis. Sequences derived from single-cell sorting was incorporated and the resulting files were clustered. Based on IMGT annotation, sequences were grouped based on IGHV and IGHJ gene identity, HCDR3 length, and nucleic acid homology across the VDJ junction. Lineage assignments were incorporated with IMGT annotations to generate an SQL database subjected to further analysis. Intra-sample clonal overlap was determined using scripts in Matlab^24^ (R2020a, The Mathworks Inc.) and in R (R core team). Phylogenetics were reconstructed using Phylip’s DNA/AA parsimony (dnapars, propars) tool (v3.695; adjusting settings 1, 4, 5, 6 and O setting the germline sequence as the outgroup. The resulting output was visualized using Cytoscape v3.8.2 to generate lineage trees.^21^

### Single cell RNA Sequencing (scRNA-Seq)

We used 5’ gene expression (5’GEX) and paired V(D)J sequencing kits from the 10X Genomics Chromium platform during scRNA-Seq analysis of FACS-sorted B cell populations. All scRNA-Seq library preparation was supported by the infrastructure and staff of the UAB Flow Cytometry and Single Cell Core and followed manufacturers guidelines. Raw sequencing data was collected on an Illumina MiSeq in the UAB Heflin Genomics Core and was processed by joining of paired end sequencing reads and filtering of low-quality sequences (Q<30), prior to being aligned to the GRCh38 reference human genome for annotation using the CellRanger Pipeline (10X Genomics, v3.1.0) hosted on the UAB Cheaha supercomputing cluster.

This produced 4 individual sequencing libraries (CD19+ and GAC+ B cell populations from two tissue donors), which were independently filtered to high quality cells (<5% mitochondrial gene content, read count 200-3000) using in Seurat Version 4.^47^ Normalized gene count matrices were derived from the top 3000 variably expressed genes. To prevent undesirable clustering of cells based on expression of unique B cell receptors, we filtered immunoglobulin-related genes from the variable gene lists for each specimen, as has been done previously.^33^ Additionally, T cells, identified by high expression of CD3D and CD3E (0.27% of all filtered cells), were removed. This trimmed the variable feature count for each CD19+ and GAC+ B cell library to 2,838 and 2,880 for A051a and 2,829 and 2,877 for A103a, respectively. We used these variable gene lists to identify integration anchors between individual libraries and integrate the samples, thereby producing a sample of 10,571 pooled cells for further analysis.

Differentially expressed transcripts used to assign cellular identify of clusters included: ASC: XBP-1 and JCHAIN, naïve B cells: S1PR1, FCER2 and CXCR4, MZ-like B cells: NOTCH2, Complement Receptor 2 (CR2, CD21/35), ZEB2. AtypBMem: FOS, Jun, IL-6, RIG-1, MDA5. B_Act_: NFkB, MYC, CD69 CD44, HLA-associated molecules. BMem: NFRSF13B, FCRL4. Pre-GC B cells: MEF2B. DZ-GCB: CXCR4, MKI67, AICDA, UNG. LZ-GCB: CD79B, RELB, NR4A1, CD40, TRAF1, ICAM1, CD83, TNFSF9, TNFRSF18.

Principal Component (PC) Analysis with the integrated data set was performed over the first 50 dimensions. We performed Uniform Manifold Approximation and Projection (UMAP) using the first 30 PCs to reduce dimensionality and visualize cellular heterogeneity. To identify transcriptionally distinct cell clusters, we constructed shared nearest neighbor (SNN) graphs applied the SNN graph to the UMAP projection. We applied Clustree to inspect the effects of varying the resolution parameter on cluster assignment of individual B cells.^48^ We used a resolution of 0.8 in our SNN graphs to govern our examine gene expression analyses, which resulted in the identification of 176 transcriptionally distinct populations of cells that grouped into 4 spatially separate superclusters in UMAP space.

### UCA and Consensus sequence generation

Consensus sequences for each described lineages were generated by MSA implemented in R.^49^ UCAs sequences were generated by using the reference sequence of the gene segment alleles (V_H_, D_H_, J_H_, V_K/L_, J_K/L_) as annotated by IMGT/HighV-QUEST and reverting all mutated residues to germline configuration. Non templated nucleotides (those annotated as N additions) were not modified. Gene blocks corresponding to the UCAs or consensus sequence for each of the clones were purchased from IDT. Gene blocks were inserted into hIGM, IGK, or IGL CLIC expression vectors to produce rhIgM.

### Surface Plasmon Resonance

Localized surface plasmon resonance (L-SPR) data were collected using a 1:1 referencing system on the Nicoya Alto HT-SPR instrument according to manufacturer’s recommendations and as described.^50^ Briefly, GAS PGPS was diluting to 500ug/mL in PBS containing 0.05% Tween-20 and immobilized on an EDC/NHS-activated carboxyl sensor (Nicoya) for 20 min. The rmAbs (analytes) were diluted to 270 µM in PBS containing 0.05% Tween-20. Single-cycle kinetic analyses were performed at 25°C with five threefold analyte dilutions, with maximum concentrations of 83nM. Kinetic fitting was performed using a 1:1 Langmuir model within the Nicoya user portal. The limit of detection for the equilibrium constant (K_D_) was estimated to be 4.3 mM. For visualization and for calculations of fold-change in affinity, if a rmAb had no detectable binding by L-SPR, a K_D_ of 10 mM was substituted for graphing.

### Statistical analyses

All statistical comparisons were performed in Prism (GraphPad Software). Comparisons of group means was made using an unpaired t-test or one-way Anova. Spearman correlations were used to assess association between two nonparametric variables. Kmeans and Hierarchical clustering, heatmap plots, and other visualizations were created using R v4.2.0 (R Core Team 2022),^51^ Matlab (R2020a, The Mathworks Inc., Natick, MA), Cytoscape v3.8.2^52^, FlowJo v10.8.1 (BD, Franklin Lakes, NJ), Prism v9.5.1 (GraphPad Software, LLC., Boston, MA).

### Data Availability

Ig sequences from single-sorted GAC-reacitve B cells have been deposited in Genebank (2022 - 3038). Single-cell transcriptomic and Ig sequences have been deposited into Geo (GSE278639)

## References

1. Muthana, S.M., Xia, L., Campbell, C.T., Zhang, Y., and Gildersleeve, J.C. (2015). Competition between serum IgG, IgM, and IgA anti-glycan antibodies. PLoS One 10, e0119298. 10.1371/journal.pone.0119298.

2. Bello-Gil, D., Audebert, C., Olivera-Ardid, S., Perez-Cruz, M., Even, G., Khasbiullina, N., Gantois, N., Shilova, N., Merlin, S., Costa, C., et al. (2019). The Formation of Glycan-Specific Natural Antibodies Repertoire in GalT-KO Mice Is Determined by Gut Microbiota. Front Immunol 10, 342. 10.3389/fimmu.2019.00342.

3. Muthana, S.M., and Gildersleeve, J.C. (2016). Factors Affecting Anti-Glycan IgG and IgM Repertoires in Human Serum. Sci Rep 6, 19509. 10.1038/srep19509.

4. Hamanova, M., Chmelikova, M., Nentwich, I., Thon, V., and Lokaj, J. (2015). Anti-Gal IgM, IgA and IgG natural antibodies in childhood. Immunol Lett 164, 40–43. 10.1016/j.imlet.2015.02.001.

5. Luetscher, R.N.D., McKitrick, T.R., Gao, C., Mehta, A.Y., McQuillan, A.M., Kardish, R., Boligan, K.F., Song, X., Lu, L., Heimburg-Molinaro, J., et al. (2020). Unique repertoire of anti-carbohydrate antibodies in individual human serum. Sci Rep 10, 15436. 10.1038/s41598-020-71967-y.

6. Schneider, C., Smith, D.F., Cummings, R.D., Boligan, K.F., Hamilton, R.G., Bochner, B.S., Miescher, S., Simon, H.U., Pashov, A., Vassilev, T., and von Gunten, S. (2015). The human IgG anti-carbohydrate repertoire exhibits a universal architecture and contains specificity for microbial attachment sites. Sci Transl Med 7, 269ra261. 10.1126/scitranslmed.3010524.

7. Purohit, S., Li, T., Guan, W., Song, X., Song, J., Tian, Y., Li, L., Sharma, A., Dun, B., Mysona, D., et al. (2018). Multiplex glycan bead array for high throughput and high content analyses of glycan binding proteins. Nat Commun 9, 258. 10.1038/s41467-017-02747-y.

8. Huflejt, M.E., Vuskovic, M., Vasiliu, D., Xu, H., Obukhova, P., Shilova, N., Tuzikov, A., Galanina, O., Arun, B., Lu, K., and Bovin, N. (2009). Anti-carbohydrate antibodies of normal sera: findings, surprises and challenges. Mol Immunol 46, 3037–3049. 10.1016/j.molimm.2009.06.010.

9. Baumgarth, N., Tung, J.W., and Herzenberg, L.A. (2005). Inherent specificities in natural antibodies: a key to immune defense against pathogen invasion. Springer Semin Immunopathol 26, 347–362. 10.1007/s00281-004-0182-2.

10. Etlinger, H.M., Julius, M.H., and Heusser, C.H. (1982). Mechanism of clonal dominance in the murine anti-phosphorylcholine response. I. Relation between antibody avidity and clonal dominance. J Immunol 128, 1685–1691.

11. Etlinger, H.M., and Heusser, C.H. (1986). T15 dominance in BALB/c mice is not controlled by environmental factors. J Immunol 136, 1988–1991.

12. New, J.S., Dizon, B.L.P., Fucile, C.F., Rosenberg, A.F., Kearney, J.F., and King, R.G. (2020). Neonatal Exposure to Commensal-Bacteria-Derived Antigens Directs Polysaccharide-Specific B-1 B Cell Repertoire Development. Immunity 53, 172–186 e176. 10.1016/j.immuni.2020.06.006.

13. Khasbiullina, N.R., Shilova, N.V., Navakouski, M.E., Nokel, A.Y., Knirel, Y.A., Blixt, O., and Bovin, N.V. (2018). Repertoire of Abs primed by bacteria in gnotobiotic mice. Innate Immun 24, 180–187. 10.1177/1753425918763524.

14. Kappler, K., Restin, T., Lasanajak, Y., Smith, D.F., Bassler, D., and Hennet, T. (2020). Limited Neonatal Carbohydrate-Specific Antibody Repertoire Consecutive to Partial Prenatal Transfer of Maternal Antibodies. Front Immunol 11, 573629. 10.3389/fimmu.2020.573629.

15. Shackelford, P.G., Nelson, S.J., Palma, A.T., and Nahm, M.H. (1988). Human antibodies to group A streptococcal carbohydrate. Ontogeny, subclass restriction, and clonal diversity. J Immunol 140, 3200–3205.

16. Mitchell, R.B., Archer, S.M., Ishman, S.L., Rosenfeld, R.M., Coles, S., Finestone, S.A., Friedman, N.R., Giordano, T., Hildrew, D.M., Kim, T.W., et al. (2019). Clinical Practice Guideline: Tonsillectomy in Children (Update). Otolaryngol Head Neck Surg 160, S1–S42. 10.1177/0194599818801757.

17. Randall, T.D., Parkhouse, R.M., and Corley, R.B. (1992). J chain synthesis and secretion of hexameric IgM is differentially regulated by lipopolysaccharide and interleukin 5. Proc Natl Acad Sci U S A 89, 962–966. 10.1073/pnas.89.3.962.

18. Mekhaiel, D.N., Czajkowsky, D.M., Andersen, J.T., Shi, J., El-Faham, M., Doenhoff, M., McIntosh, R.S., Sandlie, I., He, J., Hu, J., et al. (2011). Polymeric human Fc-fusion proteins with modified effector functions. Sci Rep 1, 124. 10.1038/srep00124.

19. Zorzoli, A., Meyer, B.H., Adair, E., Torgov, V.I., Veselovsky, V.V., Danilov, L.L., Uhrin, D., and Dorfmueller, H.C. (2019). Group A, B, C, and G Streptococcus Lancefield antigen biosynthesis is initiated by a conserved alpha-d-GlcNAc-beta-1,4-l-rhamnosyltransferase. J Biol Chem 294, 15237-15256. 10.1074/jbc.RA119.009894.

20. New, J.S., Dizon, B.L.P., King, R.G., Greenspan, N.S., and Kearney, J.F. (2023). B-1 B Cell-Derived Natural Antibodies against N-Acetyl-d-Glucosamine Suppress Autoimmune Diabetes Pathogenesis. J Immunol. 10.4049/jimmunol.2300264.

21. Felsenstein, J. (2005). PHYLIP (Phylogeny Inference Package) version 3.6. Distributed by the author. Department of Genome Sciences, University of Washington, Seattle.

22. Massoni-Badosa, R., Aguilar-Fernandez, S., Nieto, J.C., Soler-Vila, P., Elosua-Bayes, M., Marchese, D., Kulis, M., Vilas-Zornoza, A., Buhler, M.M., Rashmi, S., et al. (2024). An atlas of cells in the human tonsil. Immunity 57, 379–399 e318. 10.1016/j.immuni.2024.01.006.

23. King, H.W., Orban, N., Riches, J.C., Clear, A.J., Warnes, G., Teichmann, S.A., and James, L.K. (2021). Single-cell analysis of human B cell maturation predicts how antibody class switching shapes selection dynamics. Sci Immunol 6. 10.1126/sciimmunol.abe6291.

24. Tipton, C.M., Fucile, C.F., Darce, J., Chida, A., Ichikawa, T., Gregoretti, I., Schieferl, S., Hom, J., Jenks, S., Feldman, R.J., et al. (2015). Diversity, cellular origin and autoreactivity of antibody-secreting cell population expansions in acute systemic lupus erythematosus. Nat Immunol 16, 755–765. 10.1038/ni.3175.

25. Wang, D., Wells, S.M., Stall, A.M., and Kabat, E.A. (1994). Reaction of germinal centers in the T-cell-independent response to the bacterial polysaccharide alpha(1-->6)dextran. Proc Natl Acad Sci U S A 91, 2502–2506. 10.1073/pnas.91.7.2502.

26. de Vinuesa, C.G., Cook, M.C., Ball, J., Drew, M., Sunners, Y., Cascalho, M., Wabl, M., Klaus, G.G., and MacLennan, I.C. (2000). Germinal centers without T cells. J Exp Med 191, 485–494.

27. Kin, N.W., Stefanov, E.K., Dizon, B.L., and Kearney, J.F. (2012). Antibodies generated against conserved antigens expressed by bacteria and allergen-bearing fungi suppress airway disease. J Immunol 189, 2246–2256. 10.4049/jimmunol.1200702.

28. Covens, K., Verbinnen, B., Geukens, N., Meyts, I., Schuit, F., Van Lommel, L., Jacquemin, M., and Bossuyt, X. (2013). Characterization of proposed human B-1 cells reveals pre-plasmablast phenotype. Blood 121, 5176–5183. 10.1182/blood-2012-12-471953.

29. Reynaud, C.A., and Weill, J.C. (2012). Gene profiling of CD11b(+) and CD11b(-) B1 cell subsets reveals potential cell sorting artifacts. J Exp Med 209, 433–434. 10.1084/jem.20120402.

30. Perez-Andres, M., Grosserichter-Wagener, C., Teodosio, C., van Dongen, J.J., Orfao, A., and van Zelm, M.C. (2011). The nature of circulating CD27+CD43+ B cells. J Exp Med 208, 2565–2566. 10.1084/jem.20112203.

31. Griffin, D.O., Holodick, N.E., and Rothstein, T.L. (2011). Human B1 cells in umbilical cord and adult peripheral blood express the novel phenotype CD20+ CD27+ CD43+ CD70. J Exp Med 208, 67–80. 10.1084/jem.20101499.

32. Dunn-Walters, D.K., Isaacson, P.G., and Spencer, J. (1995). Analysis of mutations in immunoglobulin heavy chain variable region genes of microdissected marginal zone (MGZ) B cells suggests that the MGZ of human spleen is a reservoir of memory B cells. J Exp Med 182, 559–566. 10.1084/jem.182.2.559.

33. Dunn-Walters, D.K., Isaacson, P.G., and Spencer, J. (1996). Sequence analysis of rearranged IgVH genes from microdissected human Peyer’s patch marginal zone B cells. Immunology 88, 618–624.

34. Zhao, Y., Uduman, M., Siu, J.H.Y., Tull, T.J., Sanderson, J.D., Wu, Y.B., Zhou, J.Q., Petrov, N., Ellis, R., Todd, K., et al. (2018). Spatiotemporal segregation of human marginal zone and memory B cell populations in lymphoid tissue. Nat Commun 9, 3857. 10.1038/s41467-018-06089-1.

35. Bemark, M., Pitcher, M.J., Dionisi, C., and Spencer, J. (2024). Gut-associated lymphoid tissue: a microbiota-driven hub of B cell immunity. Trends Immunol 45, 211–223. 10.1016/j.it.2024.01.006.

36. Weill, J.C., Weller, S., and Reynaud, C.A. (2023). B cell diversification in gut-associated lymphoid tissues: From birds to humans. J Exp Med 220. 10.1084/jem.20231501.

37. Haji-Ghassemi, O., Blackler, R.J., Martin Young, N., and Evans, S.V. (2015). Antibody recognition of carbohydrate epitopes. Glycobiology 25, 920–952. 10.1093/glycob/cwv037.

38. Soliman, C., Walduck, A.K., Yuriev, E., Richards, J.S., Cywes-Bentley, C., Pier, G.B., and Ramsland, P.A. (2018). Structural basis for antibody targeting of the broadly expressed microbial polysaccharide poly-N-acetylglucosamine. J Biol Chem 293, 5079–5089. 10.1074/jbc.RA117.001170.

39. Comer, F.I., Vosseller, K., Wells, L., Accavitti, M.A., and Hart, G.W. (2001). Characterization of a mouse monoclonal antibody specific for O-linked N-acetylglucosamine. Anal Biochem 293, 169–177. 10.1006/abio.2001.5132.

40. Snow, C.M., Senior, A., and Gerace, L. (1987). Monoclonal antibodies identify a group of nuclear pore complex glycoproteins. J Cell Biol 104, 1143–1156. 10.1083/jcb.104.5.1143.

41. De Martin, A., Stanossek, Y., Lutge, M., Cadosch, N., Onder, L., Cheng, H.W., Brandstadter, J.D., Maillard, I., Stoeckli, S.J., Pikor, N.B., and Ludewig, B. (2023). PI16(+) reticular cells in human palatine tonsils govern T cell activity in distinct subepithelial niches. Nat Immunol 24, 1138–1148. 10.1038/s41590-023-01502-4.

42. Rosenberg, A.F., Killian, J.T., Green, T.J., Akther, J., Hossain, E., Qiu, S., Randall, T.D., Lund, F.E., and King, R.G. (2023). A high-throughput multiplex array for antigen-specific serology with automated analysis. bioRxiv, 2023.2003.2029.534777. 10.1101/2023.03.29.534777.

43. Mc, C.M., and Lancefield, R.C. (1955). Variation in the group-specific carbohydrate of group A streptococci. I. Immunochemical studies on the carbohydrates of variant strains. J Exp Med 102, 11–28. 10.1084/jem.102.1.11.

44. Nellore, A., Zumaquero, E., Scharer, C.D., Fucile, C.F., Tipton, C.M., King, R.G., Mi, T., Mousseau, B., Bradley, J.E., Zhou, F., et al. (2023). A transcriptionally distinct subset of influenza-specific effector memory B cells predicts long-lived antibody responses to vaccination in humans. Immunity. 10.1016/j.immuni.2023.03.001.

45. Tom, R., Bisson, L., and Durocher, Y. (2008). Transfection of HEK293-EBNA1 Cells in Suspension with Linear PEI for Production of Recombinant Proteins. CSH Protoc 2008, pdb prot4977. 10.1101/pdb.prot4977.

46. Brochet, X., Lefranc, M.P., and Giudicelli, V. (2008). IMGT/V-QUEST: the highly customized and integrated system for IG and TR standardized V-J and V-D-J sequence analysis. Nucleic Acids Res 36, W503–508. 10.1093/nar/gkn316.

47. Stuart, T., Butler, A., Hoffman, P., Hafemeister, C., Papalexi, E., Mauck, W.M., 3rd, Hao, Y., Stoeckius, M., Smibert, P., and Satija, R. (2019). Comprehensive Integration of Single-Cell Data. Cell 177, 1888-1902 e1821. 10.1016/j.cell.2019.05.031.

48. Zappia, L., and Oshlack, A. (2018). Clustering trees: a visualization for evaluating clusterings at multiple resolutions. Gigascience 7. 10.1093/gigascience/giy083.

49. Bodenhofer, U., Bonatesta, E., Horejs-Kainrath, C., and Hochreiter, S. (2015). msa: an R package for multiple sequence alignment. Bioinformatics 31, 3997–3999. 10.1093/bioinformatics/btv494.

50. Woodruff, M.C., Ramonell, R.P., Haddad, N.S., Anam, F.A., Rudolph, M.E., Walker, T.A., Truong, A.D., Dixit, A.N., Han, J.E., Cabrera-Mora, M., et al. (2022). Dysregulated naive B cells and de novo autoreactivity in severe COVID-19. Nature 611, 139–147. 10.1038/s41586-022-05273-0.

51. R Development Core Team (2010). R: A language and environment for statistical computing (R Foundation for Statistical Computing).

52. Shannon, P., Markiel, A., Ozier, O., Baliga, N.S., Wang, J.T., Ramage, D., Amin, N., Schwikowski, B., and Ideker, T. (2003). Cytoscape: a software environment for integrated models of biomolecular interaction networks. Genome Res 13, 2498–2504. 10.1101/gr.1239303.

