## supplemental Figures for "Human Anti-Glycan Reactivity is Driven by the Selection of B cells Utilizing Private Antibody Gene Rearrangements that are Affinity Maturated in Germinal Centers"

Figure S1.

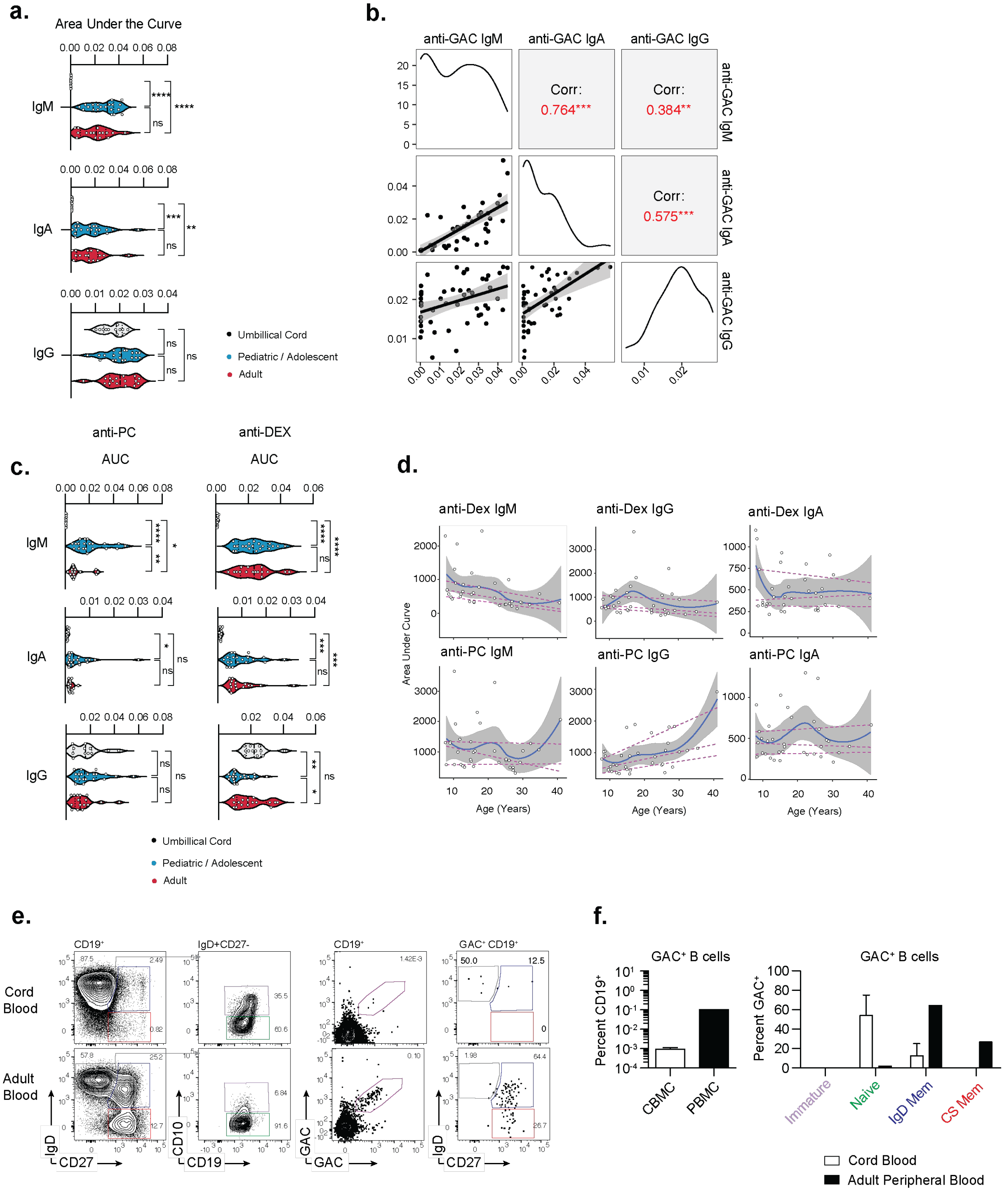


**Figure S1. Emergence of nAb reactivities and GAC-reactive B cells.**
(a) Area under the curve values of absorbance of serial dilutions of sera binding GAC measured by ELISA from the indicated age groups.

(b) Correlation area under the curve values of GAC-reactive Ab of the indicated isotypes.
(c) Area under the curve values of values of serial dilutions of sera binding the indicated antigen from the indicated age groups.
(d) Area under the curve values of Ab of the indicated isotype binding Dextran (polymers of a1-6 glucose, upper) or Phosphorylcholine (PC, lower).
(e) Representative flow cytometric histograms of B cells derived from cord blood (upper) or adult blood (lower).
(f) Frequency of $GAC^+$ B cells (left) or B cell subsets (right) from the indicated specimens.

Figure S2.

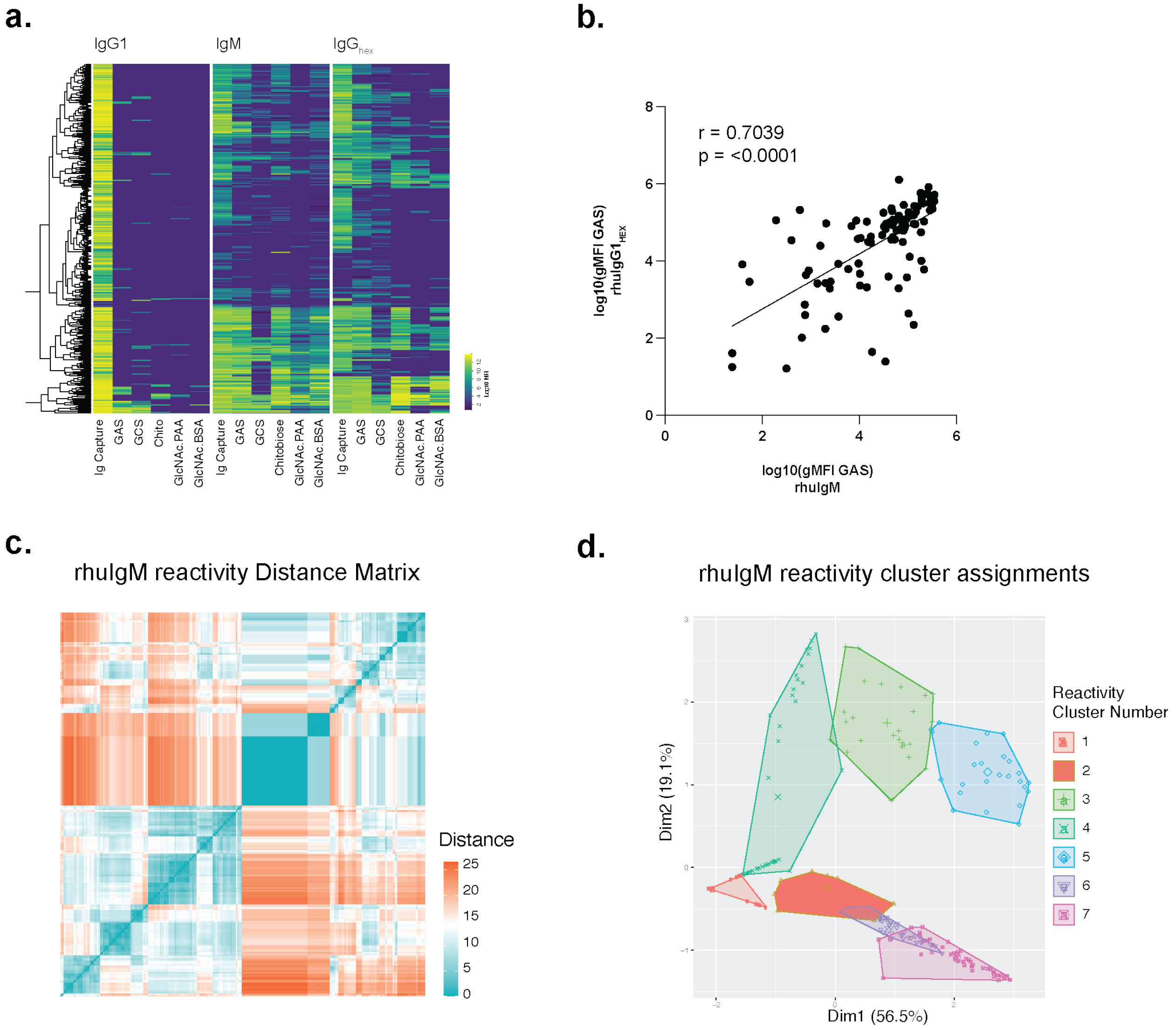


**Figure S2. GAC-reactive Abs benefit from polyvalency and display discrete reactivity profiles against GlcNAc antigens.**

**(a)** Heat map of GlcNAc array gMFI tables of a subset of tonsil GAC$^{+}$ B cell derived rAbs expressed as the indicated constructs.

**(b)** Correlation of GAC-binding rAbs expressed as IgM or IgG1$_{HEX}$.

**(c)** PCoA plot of Kmeans clustering of MFI tables of rhIgM in Figure 2.

**(d)** distance matrix of MFI tables used for reactivity-based cluster assignments.

Figure S3.

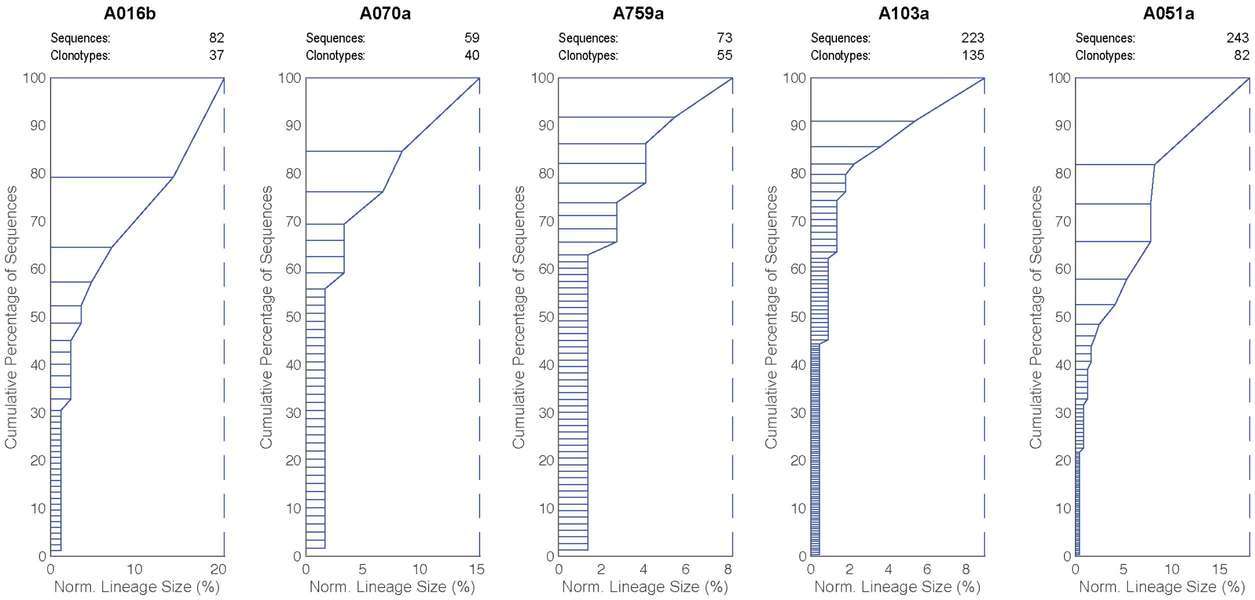


**Figure S3. GAC+ B cell clonotypes are expanded in human Tonsil.**
Clonality plots of BCR receptor sequences derived from GAC+ B cells sorted from the indicated specimens. Number of sequences recovered, lineage sizes, and cumulative percentages of each clonotype are shown.

Figure S4.

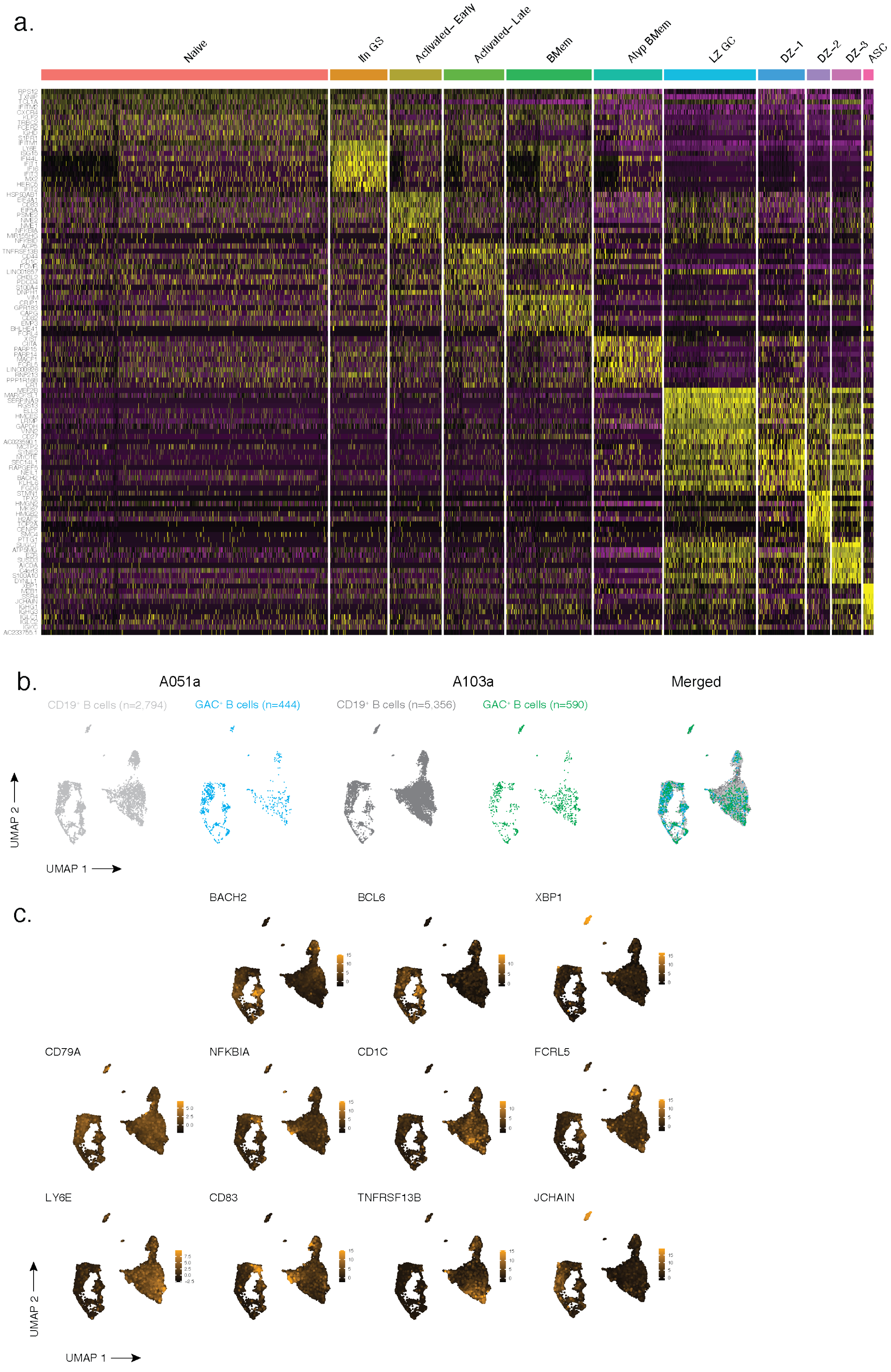
a.
Naive
Ifn GS
Activated– Early
Activated– Late
BMem
Atyp BMem
LZ GC
DZ–1
DZ–2
DZ–3
ASC
b.
A051a
A103a
Merged
CD19⁺ B cells (n=2,794)
GAC⁺ B cells (n=444)
CD19⁺ B cells (n=5,356)
GAC⁺ B cells (n=590)
UMAP 2
UMAP 1
c.
BACH2
BCL6
XBP1
CD79A
NFKBIA
CD1C
FCRL5
LY6E
CD83
TNFRSF13B
JCHAIN
UMAP 2
UMAP 1

**Figure S4. GAC$^+$ B cells are distributed through multiple B cell subsets but enriched in GC.**
**(a)** Heat map of the expression of differentially expressed genes used to annotate cluster identity.
**(b)** UMAP of CD19$^+$ and GAC$^+$ B cells defined by 5' GEX. CD19$^+$ (grey) or GAC$^+$ (colored by specimen) are indicated.
**(c)** Feature plots of indicated genes in UMAP space.

### Figure S5.

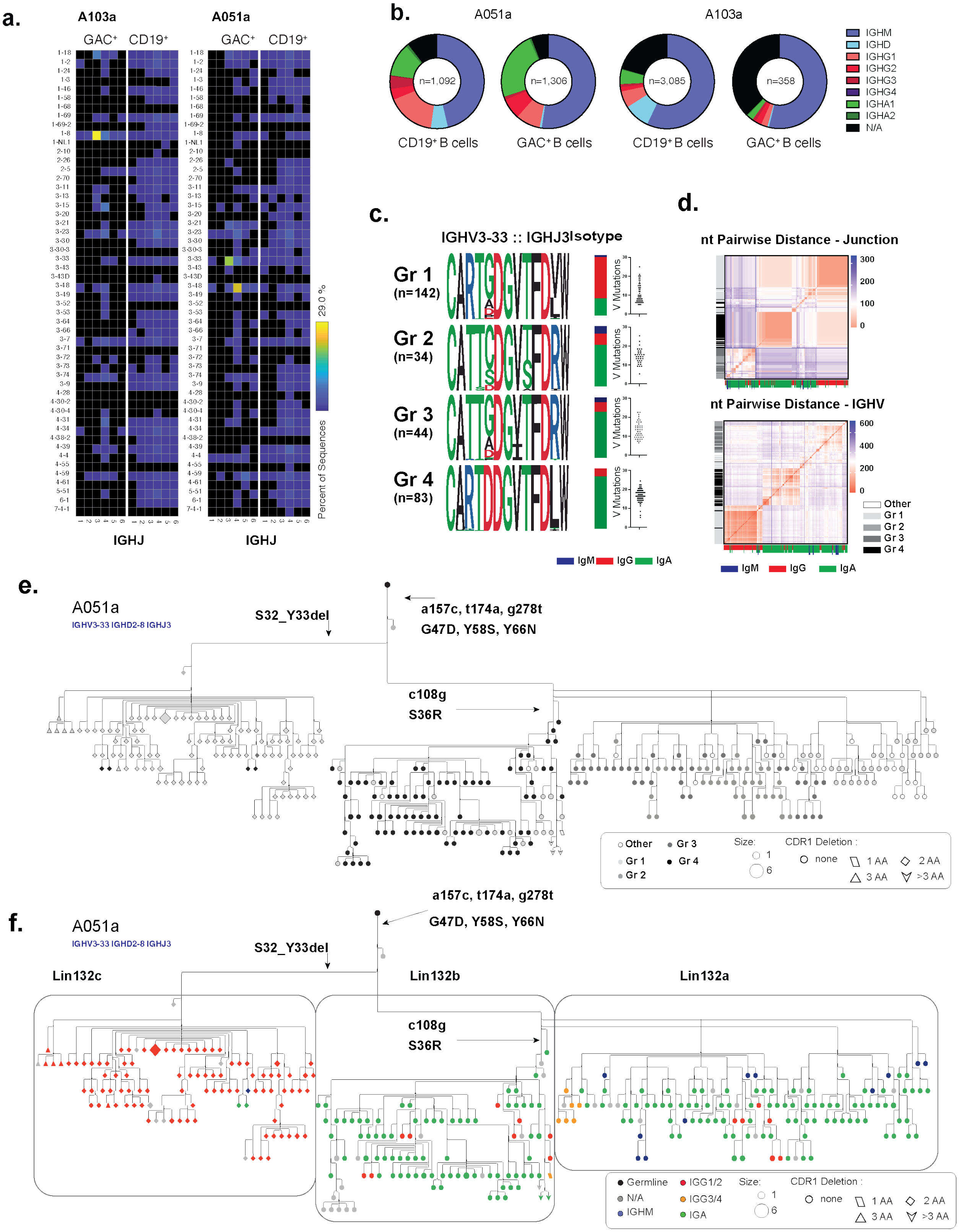

a.
A103a
A051a
GAC+ CD19+
IGHJ
Percent of Sequences
29.0 %
b.
A051a
A103a
n=1,092
n=1,306
n=3,085
n=358
CD19+ B cells
GAC+ B cells
IGHM
IGHD
IGHG1
IGHG2
IGHG3
IGHG4
IGHA1
IGHA2
N/A
c.
IGHV3-33 :: IGHJ3
Isotype
Gr 1 (n=142)
Gr 2 (n=34)
Gr 3 (n=44)
Gr 4 (n=83)
V Mutations
IgM IgG IgA
d.
nt Pairwise Distance - Junction
nt Pairwise Distance - IGHV
Other
Gr 1
Gr 2
Gr 3
Gr 4
IgM IgG IgA
e.
A051a
IGHV3-33 IGHD2-8 IGHJ3
S32_Y33del
a157c, t174a, g278t
G47D, Y58S, Y66N
c108g
S36R
Other
Gr 1
Gr 2
Gr 3
Gr 4
Size:
CDR1 Deletion :
none
1 AA
2 AA
3 AA
>3 AA
f.
A051a
IGHV3-33 IGHD2-8 IGHJ3
S32_Y33del
a157c, t174a, g278t
G47D, Y58S, Y66N
Lin132c
Lin132b
Lin132a
c108g
S36R
Germline
N/A
IGHM
IGG1/2
IGG3/4
IGA
Size:
CDR1 Deletion :
none
1 AA
2 AA
3 AA
>3 AA

**Figure S5. GAC$^+$ B cells are expanded and diversified in tonsil.**
**(a)** Heat map of IGHV and IGHJ gene usage of GAC$^+$ B cell receptors of the 10x VDJ sequences derived from the indicated tonsil specimens.
**(b)** Donut plots of the Ig isotype distribution within IGHV sequences from the indicated sorted populations Isotypes are depicted in by the indicated colors.
**(c)** Amino acid sequence logos of IGH VDJ junctions of four clonotypes identified with of all A051a derived IGHV3-33:IGHJ3 clonotypes with a junction length of 39 nt. Isotype distribution (bar plots) and somatic mutation rates (scatter plots) of members of the indicated groups are shown.
**(d)** Distance matrix heat maps of the pairwise junction nt (top) or IGH region (bottom). Lineage group assignments (x axis) and isotype (y axis) are depicted colored bars are shown.
**(e)** Phylogenic tree of nt parsimony of all members of groups 1-4. Somatic mutations corresponding to early branch points are indicated. Each sequence is colored in grey scale corresponding to its group membership. Shapes indicate number of AA deletions in CDR1 of each sequence.
**(f)** Phylogenic tree of nt parsimony of all members of groups 1-4. Sequences are colored by originating isotype.

Figure S6.

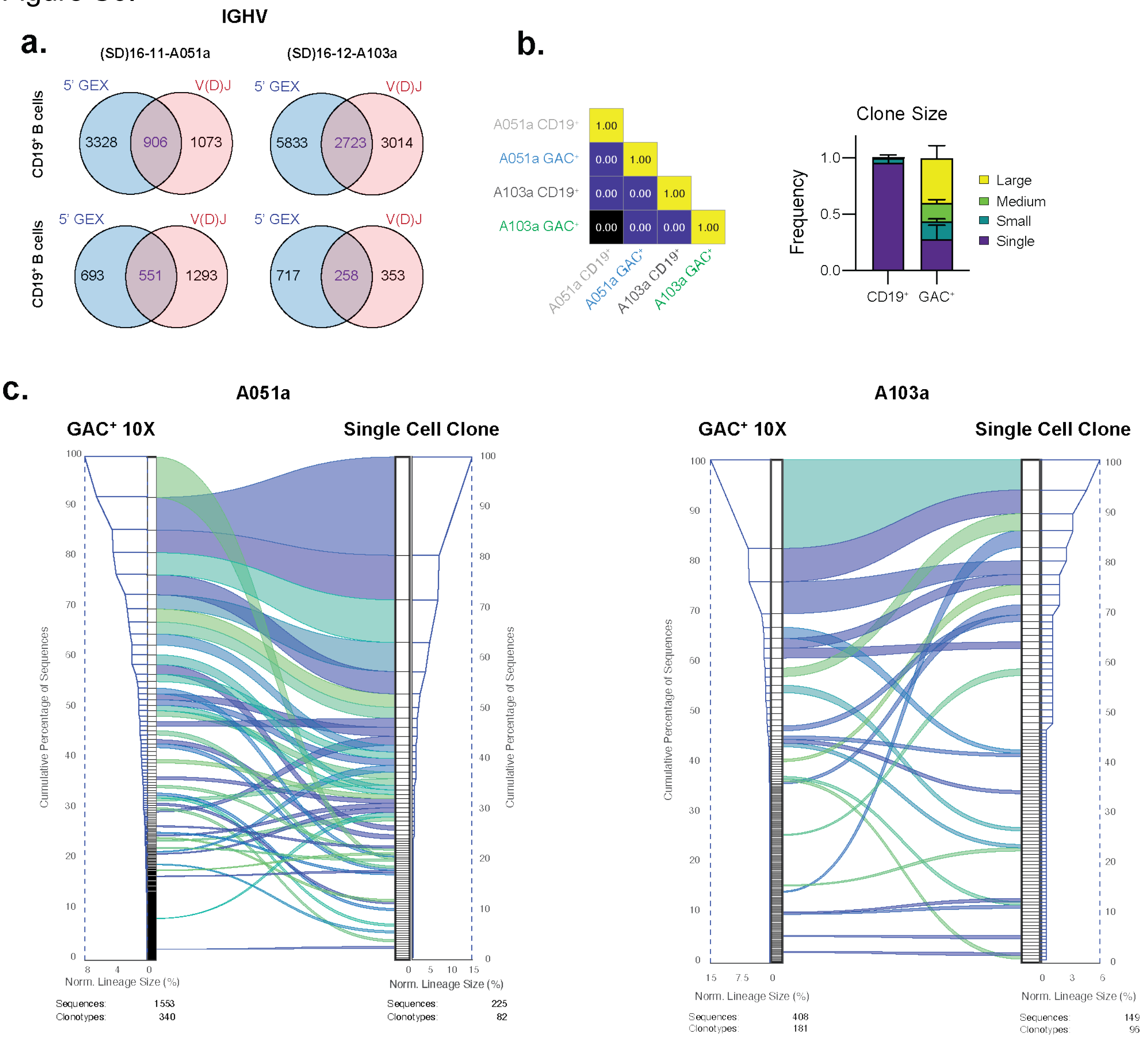


**Figure S6. Concordance between single-cell and 10x VDJ sequence recovery.**
**(a)** Venn diagrams of the filtered 10x data base. The number of cell barcodes isolated in 5' GEX and VDJ sequencing libraries and overlapping cells are shown for each sample.
**(b)** Heat map of the Morissita-Horn overlap index for each population of both libraries.
**(c)** Clonality ribbon diagrams of lineages recovered from GAC+ cells in 10x VDJ libraries (left) and single cell sorting and sequencing (right). Cumulative percentages on outer clonality plots indicate percentage of total sequences of each clone. Ribbons connect shared lineages. The number of sequences and unique lineage clonotypes are shown.

Figure S7.

a.

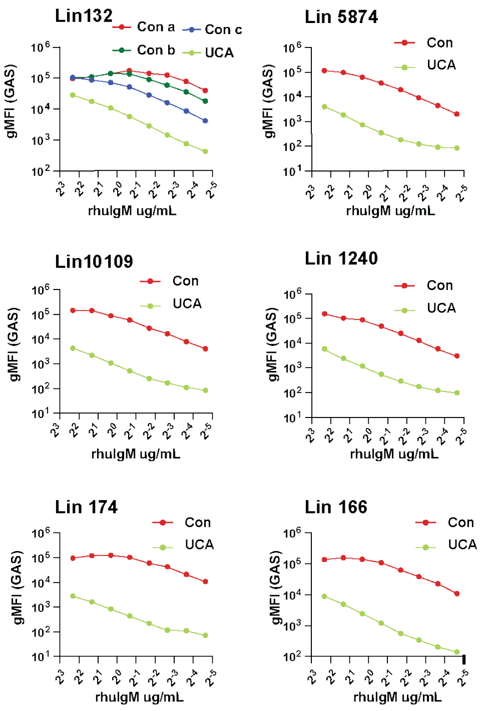


b.

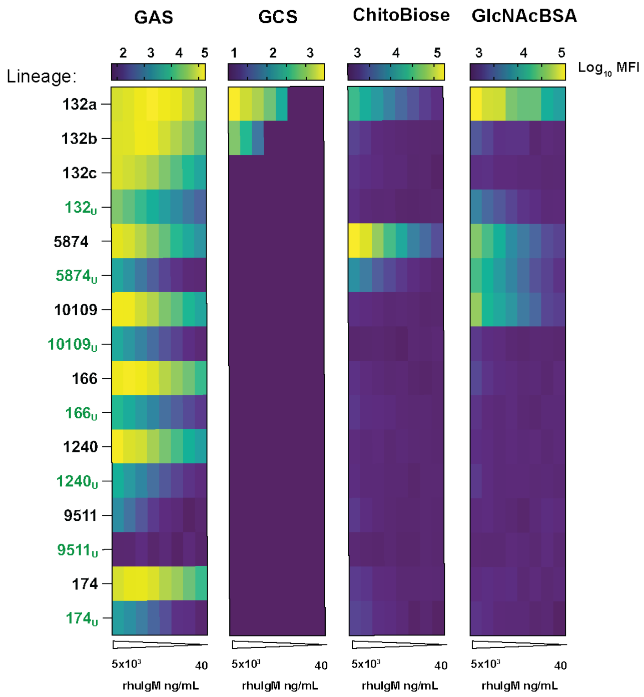


c.

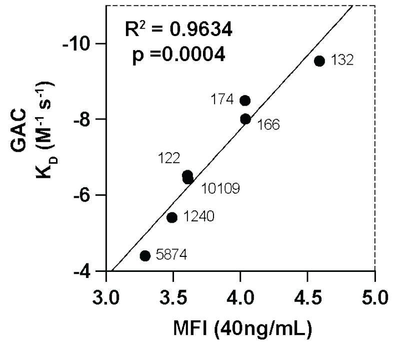


**Figure S7. Expanded GAC$^{+}$ B cells expanded B cell lineages are derived from GAC$^{+}$-reactive precursors and display heterogenous binding to GlcNAc.**
**(a)** Dilution curves of the MFI resulting from staining GAS with the rhIGM derived from the indicated lineage consensus (red) or UCA (green) at the indicated dilutions.
**(b)** Heatmap of the MFI resulting from staining of the indicated bioparticle/antigen coated beads at the indicated concentration. rhIgM derived from lineage consensus sequence are labeled in black, UCAs denoted by subscript U in green.
**(c)** Correlation of the SPR generated $K_{Dapp}$ of the indicated rhIgM and the MFI of GAS stained at 40ng/mL.
